# Rapid compensatory evolution by secondary perturbation of a primary disrupted transcriptional network

**DOI:** 10.1101/2022.06.15.496250

**Authors:** Po-Chen Hsu, Yu-Hsuan Cheng, Chia-Wei Liao, Yu-Ting Jhou, Florica Jean Ganaden Opoc, Ahmed A A Amine, Jun-Yi Leu

## Abstract

The discrete steps of transcriptional rewiring have been proposed to occur neutrally to ensure steady gene expression under stabilizing selection over long time-scales, especially when a regulon is being transferred from one transcription factor (TF) to another. Cooperative DNA binding between redundant regulatory components at the intermediate transition stage is believed to mediate this process, enabling a conflict-free switch between two TFs without a disruptive change in gene expression. Here, we have performed an evolutionary repair experiment on the *Lachancea kluyveri* yeast *sef1*Δ mutant by means of a suppressor development strategy. Complete loss of *SEF1* forced cells to activate a rewiring process to compensate for the pleiotropic defects arising from misexpression of multiple TCA cycle genes. Using different selective conditions, we identified one generalist and one specialist suppressive loss-of-function mutation of *IRA1* and *AZF1*, respectively. Our subsequent analyses show that Azf1 is a weak transcriptional activator regulated by the Ras1-PKA pathway. Azf1 loss-of-function triggers extensive gene expression changes responsible for both the compensatory and trade-off phenotypes. Our results indicate that the pleiotropic effects of dual perturbation of transcriptional networks are a potential mechanism for rapid adaptive compensation, facilitating the process of incipient transcriptional rewiring, and formation of complex traits.

## Introduction

The diversity of biological systems, including transcription regulatory systems, can be a consequence of adaptive evolution (Tenaillon et al. 2012), either macroscopically in the body plan (Peter and Davidson 2011) or microscopically at the cellular level (Lynch et al. 2014). Altered transcriptional regulation is thought to create phenotypic novelty, thereby playing a crucial role in adaptive evolution and speciation (Carroll 2000; Wagner and Lynch 2008; Romero et al. 2012; Mack and Nachman 2017). One particularly intriguing scenario is “transcriptional rewiring”, which describes changes to a gene regulatory network across species via *cis* (via polymorphisms in the linked regulatory sequences) or *trans* (through diffusible products of other loci) mechanisms over evolutionary timescales (Scannell and Wolfe 2004; Dalal and Johnson 2017). Complete regulon (a group of co-regulated and usually functionally correlated target genes) handover from one TF to another (i.e., TF substitution) is an extreme case of transcriptional rewiring (Li and Johnson 2010). The rewiring process is often constrained due to the deleterious effects of gene misexpression at intermediate stages, rendering the accumulation of regulatory changes problematic. Nevertheless, stabilizing selection in many species is believed to mitigate such constraints upon encountering new mutations by maintaining appropriate gene expression (Tanay et al. 2005; Bedford and Hartl 2009; Tirosh et al. 2009; Goncalves et al. 2012; Shi et al. 2012; Coolon et al. 2014). In other words, extensive *cis*-*trans* compensation in gene expression underlies transcriptional rewiring (Signor and Nuzhdin 2018; Signor and Nuzhdin 2019), potentially enabling “evolutionary tinkering” so that a set of conserved orthologous TFs can be fundamentally repositioned within the regulatory networks of a species (Lavoie et al. 2010). A redundancy mechanism has been proposed to explain TF substitution, whereby redundant and/or cooperative regulation underpins rewiring without giving rise to drastic changes in phenotypic output (Wohlbach et al. 2009; Johnson 2017). Interestingly, modeling of genomic data from diverse species supports that the evolution of gene expression best fits the “house-of-cards” model of stabilizing selection, whereby mutations with large effects rather than small effects evolutionarily demolish current transcriptional networks and allow them to be effectively reshuffled (Hodgins-Davis et al. 2015). However, such large-effect mutations may be a double-edged sword due to their deleterious pleiotropic effects (Dittmar et al. 2016). Therefore, we aimed to investigate this conflict, especially in terms of whether and how compensatory evolution works efficiently to deal with those pleiotropic defects as new transcriptional networks evolve, especially when the redundancy mechanism is unavailable.

To achieve this goal, we conducted an evolutionary repair experiment. Evolutionary repair is a category of experimental microbial evolution whereby the founder strain contains at least one genetic perturbation (e.g., a single gene deletion) (LaBar et al. 2020). A general concern for evolutionary repair experiments is whether such a theoretically rare event can in fact happen in nature because a genetic perturbation would be selected against in natural populations and thus kept at a low frequency. Therefore, we chose a target gene whose loss of function causes condition-dependent deleterious effects, which generally allows the perturbed founders to persist in a changing environment long enough to evolve a compensatory mutation.

Using the yeast *L. kluyveri* as a model, we first deleted the TF gene *SEF1* involved in respiration and growth (Hsu et al. 2021) to create a less-fit founder strain displaying pleiotropic defects (i.e., misregulation of multiple TCA cycle genes that affect respiration-related and -unrelated traits). Complete loss of Sef1 excludes the possibility of “stabilizing” evolution via the redundancy mechanism. Then, we performed our evolutionary repair experiment through *sef1*Δ suppressor development to screen for potential large-effect compensatory mutations. We further investigated the pleiotropic effects caused by the *azf1* loss-of-function mutation. Our results demonstrate a process of “quick-and-dirty” compensatory evolution following a sudden loss of a TF under specific conditions that precluded a smooth transition via the redundancy mechanism, potentially mimicking the extreme case of the incipient stage of transcriptional rewiring. Moreover, we also demonstrate how complex traits can easily form from perturbation of an important transcriptional network.

## Results

### Evolutionary repair rapidly suppressed the fitness defects caused by *sef1*Δ transcriptional network perturbation

Consistent with our previous work (Hsu et al. 2021), the *L. kluyveri sef1*Δ mutant is slightly less fit (slower growing) than *SEF1* strains maintained at normal temperature (28°C) under fermentative conditions [YPD: Yeast extract-Peptone-Dextrose (Dex)], with incubation under respiratory conditions [YPGly: Yeast extract-Peptone-Glycerol (Gly)] or heat stress (37−39°C) further impairing growth (Fig. 1A). The diminished maximal growth rates of the *sef1*Δ mutant calculated from source growth curves (Fig. S1A and S1B) under two respiratory conditions (post-diauxic shift and YPGly, Fig. S1C) are shown in Figure 1B. An RNA-seq analysis supports these phenotypic outcomes, showing clear downregulation of multiple TCA cycle genes required for respiratory and optimal fermentative growth (Fig. 1C, Table S1, and Table S2).

**Figure 1.**
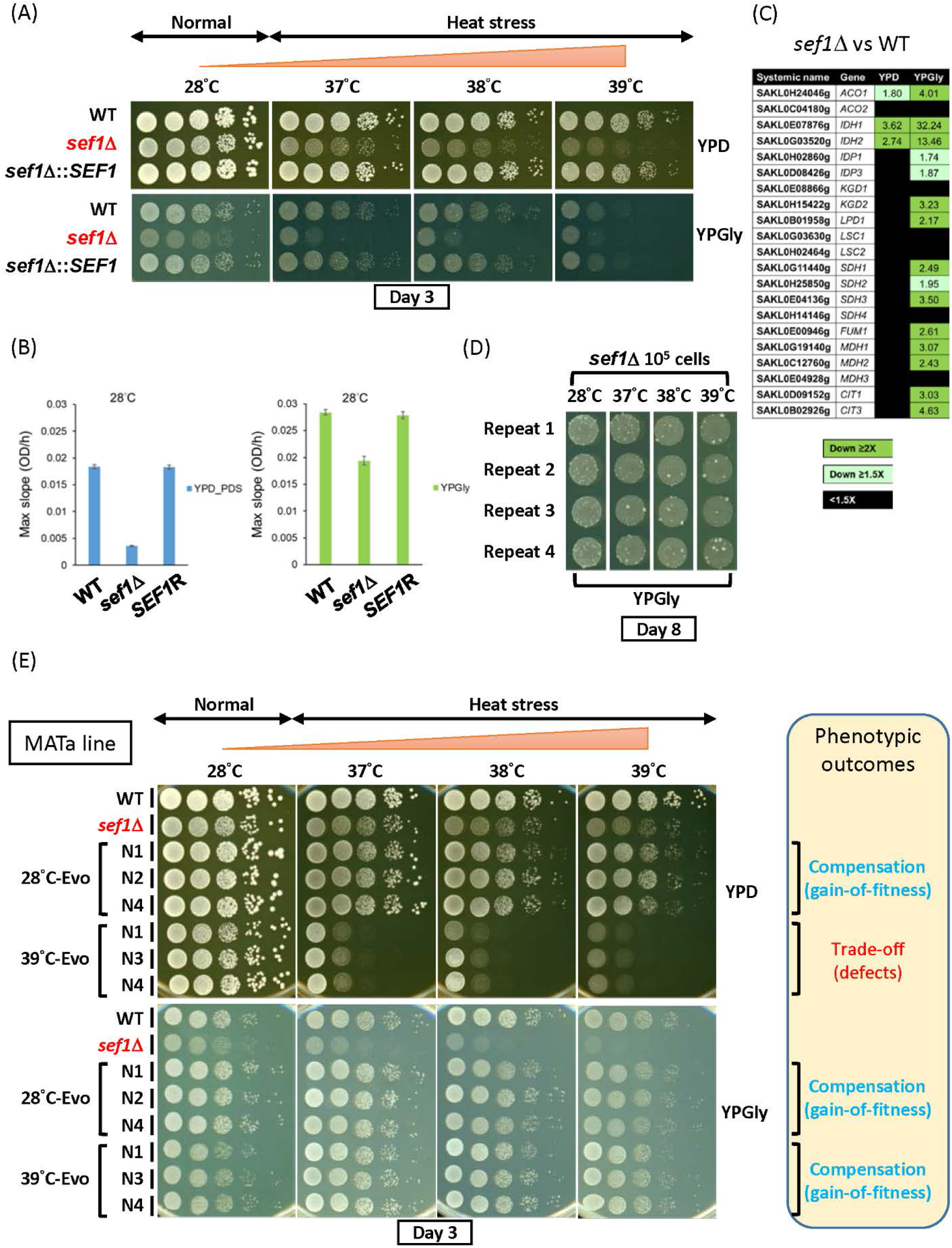
Rapid suppressor development of the *sef1*Δ mutant. (A) Growth of the *sef1*Δ mutant in response to YPD, YPGly, and heat stress. All plates were incubated for 3 days. (B) The maximal slope growth rate (OD_600_/hour) of the *sef1*Δ mutant in liquid cultures. The growth curves were measured at 28°C by using a Tecan plate reader with intermittent shaking. Error bars represent ± standard deviations. YPD_PDS: YPD culture grown to post-diauxic shift phase. (C) Differential expression of TCA cycle genes in response to *sef1*Δ. The RNA expression profiles from *L. kluyveri* wild-type and *sef1*Δ cells grown to early log-phase under both YPD and YPGly conditions are compared. A total of 21 annotated TCA cycle genes are listed, 18 of which are direct targets of Sef1 and the remaining three (*IDP1*, *MDH3*, and *CIT3*) are not (Hsu et al. 2021). (D) Rapid formation of spontaneous suppressors of *sef1*Δ phenotypes. The *sef1*Δ cells were spotted on YPGly plates at a density of 10^5^ CFU/5 μl spot and incubated at the indicated temperatures for 8 days. Cells of ancestral fitness were grown as a background lawn, while the suppressor clones were grown as bigger colonies. (E) The suppressive growth phenotypes of re-purified *sef1*Δ suppressor clones (28°C- and 39°C-Evo, MATa line).

Interestingly, we noticed that some colonies displaying greater fitness formed among the *sef1*Δ population during the growth assay, especially under the more stringent conditions (YPGly + heat stress). Increasing the incubation time accentuated the outcompeting growth of these fitter cells relative to congeners (Fig. 1D), suggesting that selection for suppressors of *sef1*Δ mutant phenotypes may occur rapidly and efficiently. Such suppressive mutations could arise from *de novo* mutations formed after selection or represent low-frequency genetic variants that pre-existed in the founder colony before selection (Teng et al. 2013). In terms of this latter, the hypothetical presence of heterogeneous quasispecies founder colonies (Fig. S1D) may accelerate the evolution of cells upon losing *SEF1*, changing their evolutionary trajectories (Fig. S1E). Therefore, we performed a large-scale *sef1*Δ suppressor development experiment (in this study, we also consider it as an evolutionary repair experiment) to investigate how cells evolve to deal with the loss of the transcriptional hub *SEF1* that affects multiple downstream fitness-contributing genes.

First, we created the *sef1*Δ and then the *sef1*Δ*chs3*Δ (tetrad dissection-competent) lines from the wild-type MATa strain, before performing a mating-type switch to obtain a MATα strain (Fig. S2A). Two independent suppressor development experiments were performed by selecting 10^8^ MATa and MATα founder cells on YPGly plates at 28−39°C for 8 days (Fig. S2B, step A). During that process, larger suppressor colonies carrying the correct drug markers were picked (Fig. S2C, steps A, B, and C). Finally, a total of 240 *sef1*Δ suppressors were isolated, including 144 MATa lines and 96 MATα lines that evolved at different temperatures (Fig. S3A). To characterize the suppressive (adaptive) effects of each clone, we evaluated their fitness by comparing their growth patterns on agar plates against two reference strains (the wild type and the corresponding *sef1*Δ founder), and then assigned a simple fitness score under YPD and YPGly conditions based on fitness categories (Fig. S3B, left and middle columns, with two examples shown in Fig. S3C). The simple fitness scores of all 240 clones are shown in Table S3. To visualize global fitness patterns among the different groups of suppressor clones, we averaged the simple fitness scores and highlighted them with different colors according to the range of mean scores (Fig. S3B, right column, and summarized in Fig. S3D and S3E). In general, the *sef1*Δ suppressor selected at 28°C (hereafter, 28°C-Evo) displayed more improved fitness than the *sef1*Δ founder under both YPD and YPGly conditions, irrespective of temperature. Hereafter, we describe the 28°C-Evo clones as “double-compensation”. In contrast, the *sef1*Δ suppressors selected under heat-stressed conditions (hereafter, 37°C-, 38°C-, or 39°C-Evo) showed more improved fitness compared to the *sef1*Δ founder only under YPGly conditions and especially under higher temperatures. Moreover, the 37, 38, and 39°C-Evo clones showed a severe growth defect relative to the *sef1*Δ founder when grown under heat-stressed YPD conditions. We specifically describe these heat-stressed 37°C-, 38°C-, and 39°C-Evo clones as “Dex-trade-off and Gly-compensation” hereafter. Only 30 of the 240 (12.5%) suppressor clones showed inconsistent phenotypic patterns (Fig. S3F), indicating that both the “double-compensation” and “Dex-trade-off and Gly-compensation” phenotypes are quite representative, implying that at least two major trajectories underlie the rapid evolutionary repair response to *SEF1* deletion.

### Suppressive phenotypes are monogenically induced by loss-of-function *azf1* or *ira1* mutations

To identify the causal mutations giving rise to the evolved phenotypes, we repurified the suppressor clones from −80°C stocks and confirmed both the “double-compensation” and “Dex-trade-off and Gly-compensation” phenotypes for two independent MATa and MATα lines (Fig. 1E and S4A). Notably, the randomly picked 37°C-Evo and 38°C-Evo clones showed the same pattern of evolved phenotypes as the 39°C-Evo clones did (Fig. S4B), implying that they may carry mutations eliciting similar suppressive mechanisms. Therefore, only three clones each of 28°C-Evo and 39°C-Evo were re-stocked, denoted “N” as indicated in Figure 1E and Figure S4A, and then subjected to whole-genome sequencing.

By comparing the resulting genomes with those of their ancestors (i.e., the founder strains), we identified mutated loci in all three clones from the two independent lines (MATa and MATα) (Table S4, Fig. 2A, and 2B). Interestingly, only two (*IRA1* and *FLO1*) and four (*FLO1*, *AZF1*, SAKL0D15356pg, *AMD2*) loci were shared among the respective 28°C-Evo and 39°C-Evo lines. The allele frequencies of the *FLO1* and *AMD2* mutations are <100% and quite diverse between clones, and SAKL0D15356pg is a pseudogene. Subsequent examination of mutation types revealed that only the *IRA1* and *AZF1* loci carry potential (missense mutations) and absolute loss-of-function mutations (premature stop codon gained or frameshift) (Fig. 2C and 2D), supporting that *ira1* and *azf1* loss-of-function mutations are causal in the 28°C-Evo and 39°C-Evo suppressors, respectively.

**Figure 2.**
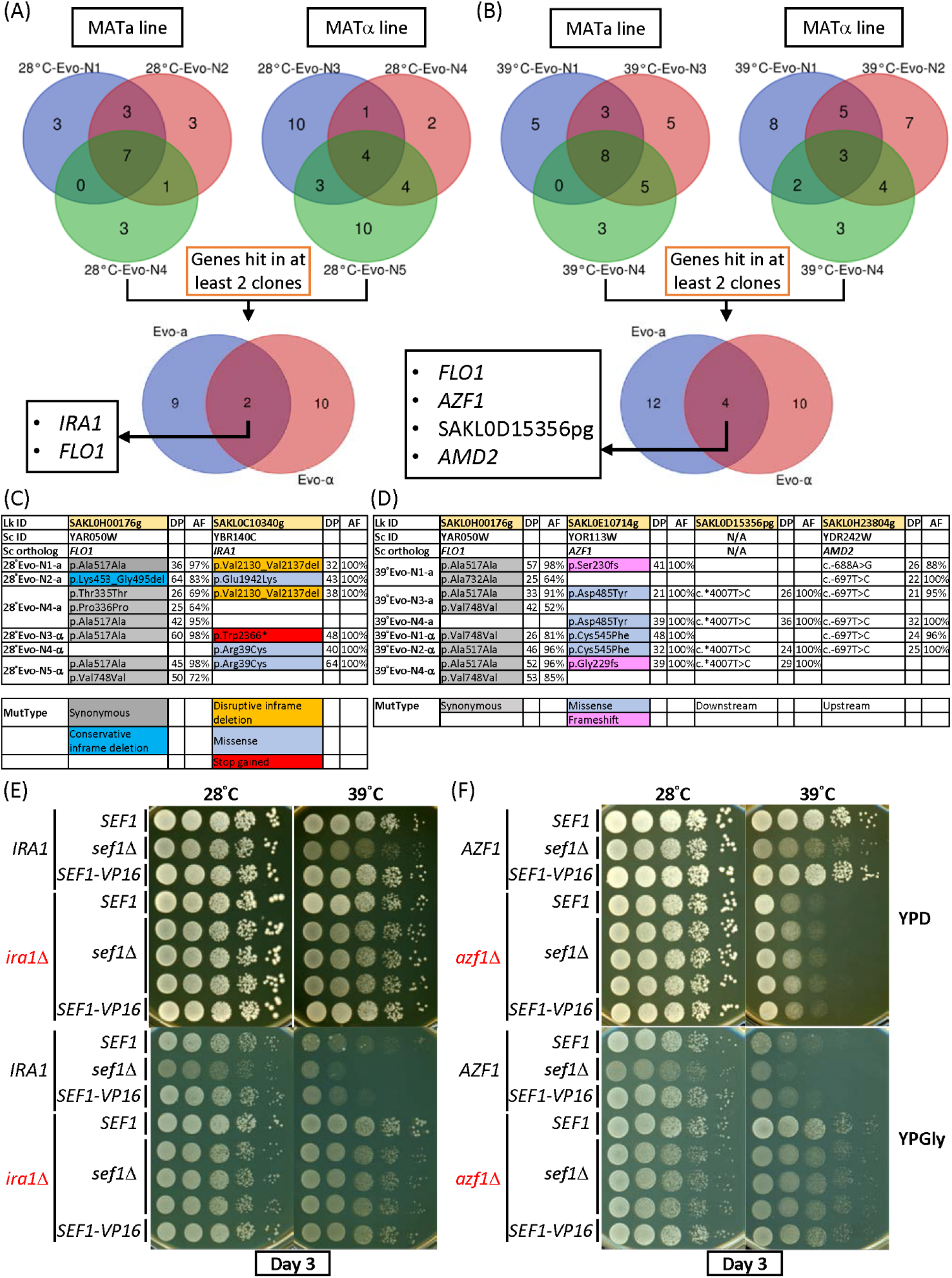
Identification of causal suppressive mutations of *sef1*Δ phenotypes. **(A)** Convergent mutations among 28°C-Evo *sef1*Δ suppressors between the MATa and MATα lines. (B) Convergent mutations among 39°C-Evo *sef1*Δ suppressors between the MATa and MATα lines. Types of candidate causal mutations in (C) 28°C-Evo *sef1*Δ suppressors and (D) 39°C-Evo *sef1*Δ suppressors. DP: read depth; AF: allele frequency. The mutant types (MutType) are highlighted with different background colors. (E) The suppressive growth phenotypes of re-constructed *sef1*Δ*ira1*Δ strains. (F) The suppressive growth phenotypes of re-constructed *sef1*Δ*azf1*Δ strains. Three independent *sef1*Δ*ira1*Δ and *sef1*Δ*azf1*Δ clones were tested. *SEF1-VP16* represents the high-activity *SEF1* strain created by fusing the strong viral transcriptional activator VP16 to the C-terminal of *SEF1* as described (Hsu et al. 2021), and one clone for each scenario is shown.

To prove that the suppressive phenotypes are monogenic, we performed tetrad dissection analyses by backcrossing the suppressor clones with their founders and then checking three MATa 28°C-Evo and MATα 39°C-Evo clones. After sporulation, all four spores of each tetrad were phenotyped and the candidate causal mutation loci were sequenced. All tetrads showed a perfect 2-to-2 ratio between suppressive vs wild-type phenotypes, consistent with the 2-to-2 genotypes (Fig. S5 and S6), indicating a clear monogenic effect of the suppressive mutation.

As expected, deletion of *IRA1* or *AZF1* alone in the parental *sef1*Δ strains proved sufficient to phenocopy the 28°C-Evo and 39°C-Evo suppressors, respectively (Fig. 2E and 2F). Interestingly, the “double-compensation” effect of *ira1*Δ and the “Dex-trade-off and Gly-compensation” effect of *azf1*Δ were both retained in the wild-type *SEF1* and high-activity *SEF1* (Hsu et al. 2021) backgrounds, indicating that these two genes can function independently of and epistatic to *SEF1*. Given general concepts of “modular epistasis” among members in an interaction network (Segrè et al. 2005) and that suppressive interactions (query and suppressive mutations) tend to connect functionally related genes (van Leeuwen et al. 2016), we hypothesized that Sef1, Ira1, and Azf1 may be associated with related functional modules.

### Sef1, Ira1, and Azf1 belong to related functional modules

*L. kluyveri IRA1* is homologous to a negative regulator of the Ras signaling pathway in *Saccharomyces cerevisiae* (Tanaka et al. 1990), which is generally coupled with the cAMP-PKA pathway that is activated in response to nutrients and the environment and is responsible for regulating metabolism, cell growth, stress resistance, and glucose adaptation (Conrad et al. 2014). *L. kluyveri* Sef1 is a transcriptional activator (Hsu et al. 2021), whereas Azf1 is a predicted TF based on orthology to *S. cerevisiae* Azf1 (Stein et al. 1998; Slattery et al. 2006). We speculated that Ras1-Ira1-cAMP-PKA represents the regulatory signaling pathway upstream of Sef1 and Azf1, so we used either chromosome-integrated or plasmid-based one-hybrid assays (Fig. 3A) to evaluate the transcriptional activity of Sef1 and Azf1 in response to Ras1-Ira1-cAMP-PKA pathway signaling. We adopted five Ras1-Ira1-cAMP-PKA hyperactive mutants for this experiment, namely *ira1*Δ (loss of a Ras1 negative regulator), *RAS1*^G20V^ (a hyperactive GTP-bound Ras1 mutant orthologous to *S. cerevisiae RAS2*^G19V^)(Toda et al. 1985), *pde2*Δ and *pde1*Δ*pde2*Δ (major loss-of-function mutants of high-affinity cyclic AMP phosphodiesterase), and *bcy1*Δ (loss of the cAMP-dependent protein kinase A inhibitor subunit). As shown in Figure 3B and 3C, hyperactivation of the Ras1-Ira1-cAMP-PKA pathway modulated Sef1 activity by repressing it under the YPD condition and activating it under the YPGly condition. The changes in Sef1 protein abundance in response to *ira1*Δ were consistent with altered activity (Fig. 3D), implying that the underlying regulatory mechanism operates via protein expression or stability.

**Figure 3.**
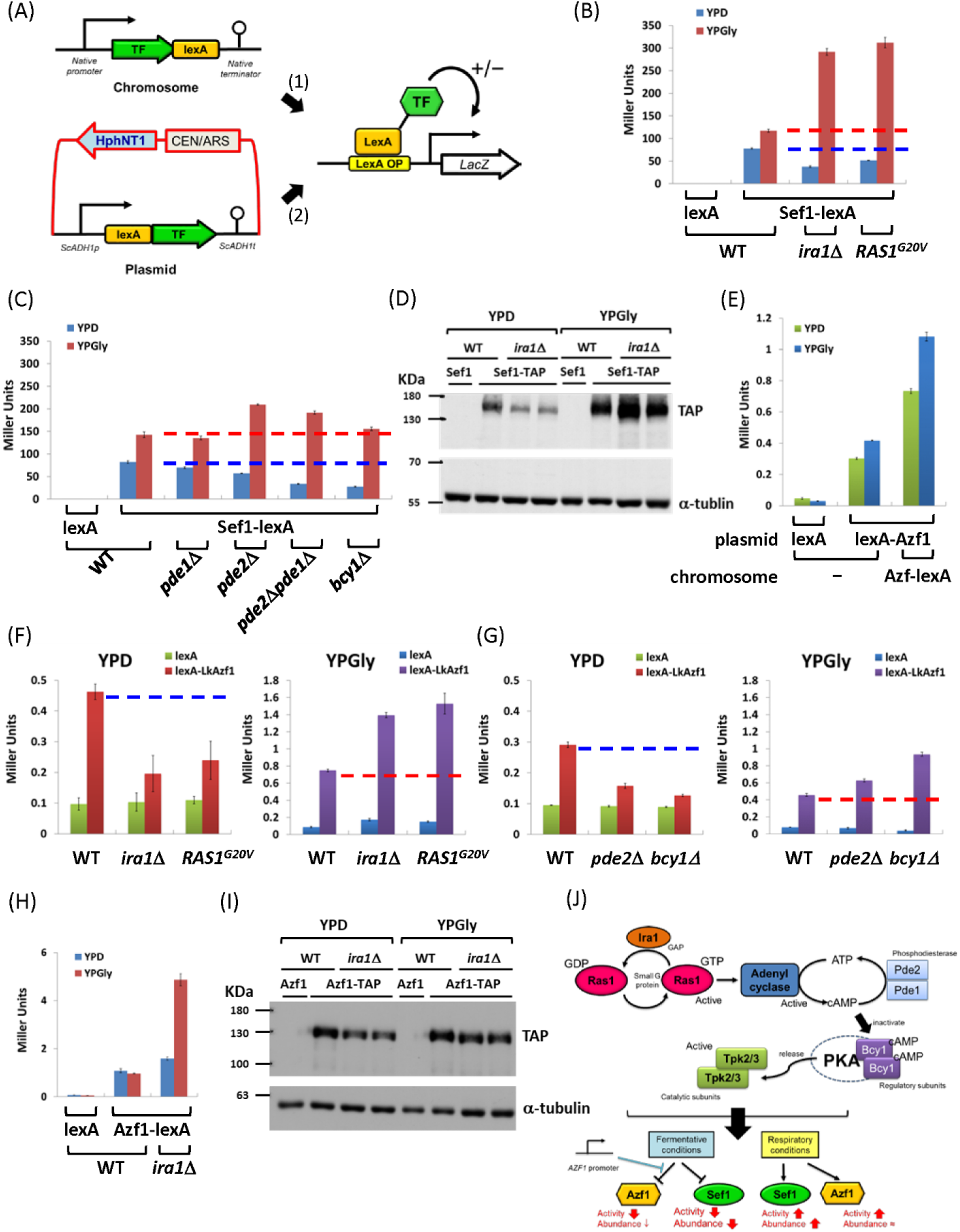
The Ras1-Ira1-PKA pathway regulates Sef1 and Azf1. (A) Two one-hybrid systems for evaluating transcriptional activities of a TF to the LacZ reporter (Hsu et al. 2021). The first is the chromosome-integrated system in which the target is C-terminally fused to the lexA domain and then expressed at the native locus. The second is the plasmid-based system, whereby the target is N-terminally fused with the lexA domain and is then expressed under the control of a yeast constitutive promoter. (B-C) Transcriptional activity of Sef1 in response to hyperactive Ras1-Ira1 (B) and cAMP-PKA (C) pathways. (D) Protein abundance of Sef1 in response to *ira1*Δ mutation. (E) Transcriptional activity of Azf1. Both the plasmid-based and chromosome systems were used and compared to enhance detection of weak Azf1 activity. (F-G) Transcriptional activity of Azf1 in response to hyperactive Ras1-Ira1 (F) and cAMP-PKA (G) pathways. (H) Transcriptional activity of Azf1 under the control of a native promoter in response to *ira1*Δ mutation. (I) Protein abundance of Azf1 in response to *ira1*Δ mutation. (J) Summarized effects of the hyperactive Ras1-Ira1-cAMP-PKA pathway on Sef1 and Azf1. For (B), (C), (E), (F), (G), and (H), LacZ activity was measured using liquid-galactosidase assay, and results are displayed as average Miller units ± SD from at least three technical repeats.

Next, we investigated Azf1 according to the same strategy described above, which demonstrated that *L. kluyveri* Azf1 is a weak transcriptional activator (Fig. 3E). Interestingly, hyperactivation of the Ras1-Ira1-cAMP-PKA pathway modulated Azf1 activity in the same way as observed for Sef1 (Fig. 3F and 3G). However, repression of Azf1 activity under the YPD condition was not detected when we used the chromosome-integrated one-hybrid constructs (i.e., in which the native promoter was used to express Azf1 fused with C-terminal lexA) (Fig. 3H), and altered Azf1 activity could not be explained by changes in protein abundance (Fig. 3I). These results imply that Azf1 is regulated by a different and more complex Ras1-Ira1-cAMP-PKA pathway-mediated mechanism than Sef1 (Fig. 3J).

### Trade-off effects of the loss-of-function *azf1* mutation

Mutations of *IRA1* and its homologs, as well as for members of the entire Ras1-PKA pathway, have been identified previously as adaptive hotspots in numerous experimental evolution and suppressor development studies (Kao and Sherlock 2008; Parts et al. 2011; Wenger et al. 2011; Kvitek and Sherlock 2013; Lang et al. 2013; van Leeuwen et al. 2016; Venkataram et al. 2016; Huang et al. 2018; Li et al. 2018; Gorter de Vries et al. 2019; Kinsler et al. 2020; Amine et al. 2021; Johnson et al. 2021). All such mutations, which arose repeatedly, proved beneficial, globally enhancing fitness irrespective of the diverse laboratory conditions under which the experiments were conducted. Therefore, we chose to focus on the *azf1* mutation, the “Dex-trade-off and Gly-compensation” effect of which has not been characterized previously.

To identify the genes regulated by Azf1, first we confirmed that heat stress did not drastically alter the steady-state transcriptional activity of Azf1 (Fig. S7A) but only reduced the abundance of Azf1 proteins (Fig. S7B). Accordingly, deletion of *AZF1* may trigger responses similar to those observed in cells suffering heat stress. For simplicity, we decided to compare RNA expression profiles directly under the normal temperature (28°C) to reduce profile complexity as much as possible. We undertook five comparisons—*sef1*Δ/wild-type, *azf1*Δ/wild-type, *sef1*Δ*azf1*Δ/wild-type, *sef1*Δ*azf1*Δ/*azf1*Δ, and *sef1*Δ*azf1*Δ/*sef1*Δ—and differential gene expression for each is summarized in Figure S7C-S7G. Detailed profiles and gene ontology (GO) analyses are presented in Table S1, S2, and S5-S12. In general, we found that the *sef1*Δ mutation affects fewer genes (Fig. S7C and S7F) than the *azf1*Δ mutation does (Fig. S7D, S7E, and S7G).

To further link altered gene expression with *azf1*Δ-induced phenotypic outcomes, we simplified the enriched GO terms by classifying them as *sef1*Δ-driven (*sef1*Δ/wild-type and *sef1*Δ*azf1*Δ/*azf1*Δ) or *azf1*Δ-driven (*azf1*Δ/wild-type, *sef1*Δ*azf1*Δ/wild-type, and *sef1*Δ*azf1*Δ/*sef1*Δ). As shown in Fig. 4A, under the YPD condition, the *azf1*Δ-driven group displays a consistent pattern for enriched GO terms for downregulation of carbohydrate metabolic process, fatty acid metabolic process, alpha-amino acid metabolic process, and phosphorus metabolic process. In contrast, the *sef1*Δ-driven group only displays consistent enrichment for GO terms relating to downregulation of the isocitrate metabolic process and the TCA cycle. We speculated that the “Dex-trade-off” effect of *azf1*Δ may be attributable to this downregulation. The GO term “carbohydrate metabolic process” of the *azf1*Δ-driven group encompasses 80 downregulated genes, including those about core glucose utilization pathways and especially glycolysis (Fig. S8), indicating that glycolysis in the *azf1*Δ cells may be particularly vulnerable. Indeed, both the *azf1*Δ and *sef1*Δ*azf1*Δ mutant lines proved more sensitive to the glycolysis inhibitor, 2-deoxyglucose (Laussel and Léon 2020), especially at higher temperatures (Fig. 4B, 4C, S9A, and S9B).

**Figure 4.**
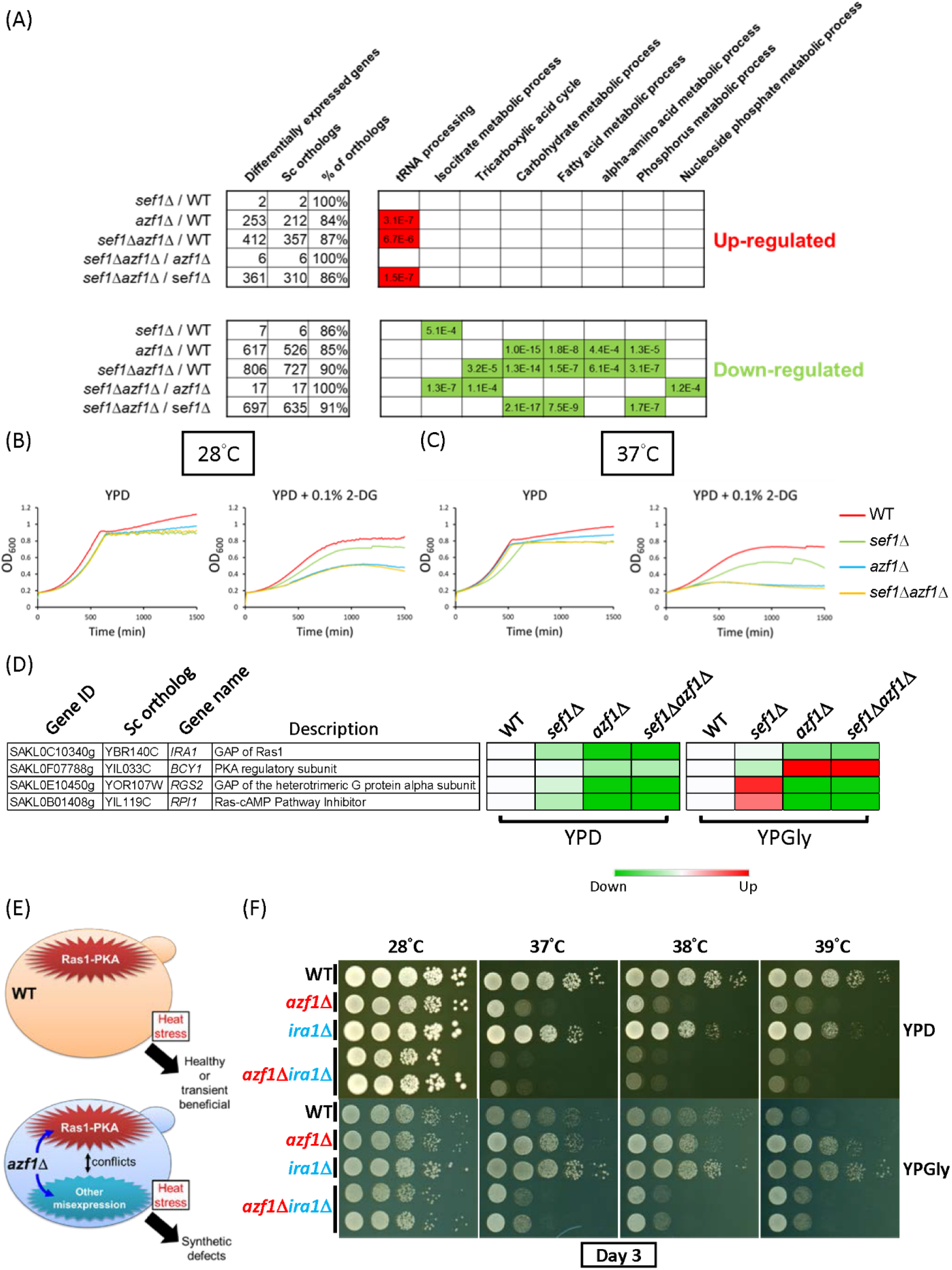
Phenotypic response to *azf1*Δ mutation in the YPD condition. (A) Selected GO terms enriched under the YPD condition. The adjusted *p*-values of the hypergeometric test with Bonferroni correction are shown. (B-C) Hypersensitivity of *azf1*Δ mutants to 2-deoxyglucose under the YPD condition without heat stress (28°C) (B) or with mild heat stress (37°C) (C). (D) The major negative regulators of the Ras1-PKA pathway are downregulated in response to *azf1*Δ mutation under the YPD condition. The heatmap was generated from the mean TPM (Transcripts Per Million) ratio of RNA-seq data relative to wild-type for each condition. (E) A proposed model describing how a hyperactive Ras1-PKA pathway due to downregulation of negative regulators (in the *azf1*Δ mutant) or *ira1*Δ mutation (used in experiments) is incompatible with the other gene misexpression induced by *azf1*Δ. (F) The synthetic growth defect of *azf1*Δ*ira1*Δ under the wild-type background in heat-stressed conditions.

Moreover, we noticed that several major negative regulators of the Ras1-PKA pathway, including the aforementioned *IRA1* and *BCY1*, were downregulated under the YPD condition (Fig. 4D). Considering that Ras1-PKA is also a glucose-responsive signaling pathway (Conrad et al. 2014), misexpression of those negative regulators may contribute to the “Dex-trade-off” effect due to conflict between the hyperactive Ras1-PKA pathway and the phenotypic impacts of other *azf1*Δ-driven gene misregulations (Fig. 4E). To test that idea, we deleted *IRA1* and *AZF1* in the wild-type or *sef1*Δ background to mimic the extreme downregulation of *IRA1*. Interestingly, both the *azf1*Δ*ira1*Δ and *sef1*Δ*ira1*Δ*azf1*Δ mutant lines showed a heat stress-induced synthetic growth defect under both YPD and YPGly conditions (Fig. 4F and S9C). Furthermore, the spores carrying both the mutated *ira1* and *azf1* alleles from tetrads of mating products between 28°C-Evo and 39°C-Evo suppressors also exhibited the synthetic growth defect (Fig. S10). These results not only support the notion that misregulation of the Ras1-PKA pathway participates in the “Dex-trade-off” effect of *azf1*Δ, but also provide corroborating evidence that Ira1 and Azf1 belong to the same functional module.

Similarly, subsequent examination of downregulated genes relating to the GO term “alpha-amino acid metabolic process” implied that multiple genes involved in Ser, Cys, Asp, Lys, Met, Gln, Glu, Pro, Arg, and Ile metabolism may be defective (Fig. S11A). This scenario is supported by the exacerbated delay in growth (longer lag-phase) of the *azf1*Δ and *sef1*Δ*azf1*Δ mutants when cells were pre-starved in an amino acid-depleted medium (Fig. S11B). Notably, we observed delayed growth at normal temperature, which was more severe under heat stress. Thus, taken together, our findings show that under the YPD condition, the *azf1*Δ mutation places cells in a state of poor core carbon and nitrogen metabolism so that their growth further deteriorates under heat stress, contributing to the “Dex-trade-off” effect.

### Beneficial effects of the loss-of-function *azf1* mutation

To connect altered gene expression with the *“*Gly-compensation” effect, we investigated the enriched GO term list under the YPGly condition (Fig. 5A). The *azf1*Δ-driven group displays a consistent pattern of enriched GO terms relating to upregulation of the cell cycle, cytoskeleton organization, DNA repair and replication, response to stress, vesicle-mediated transport, and chromatin silencing, as well as downregulation of genes related to ribosome/rRNA and tRNA metabolism that are key components in translation. In contrast, the *sef1*Δ-driven group only displays consistent enrichment for GO terms pertaining to downregulation of the TCA cycle, as reported in a previous study (Hsu et al. 2021).

**Figure 5.**
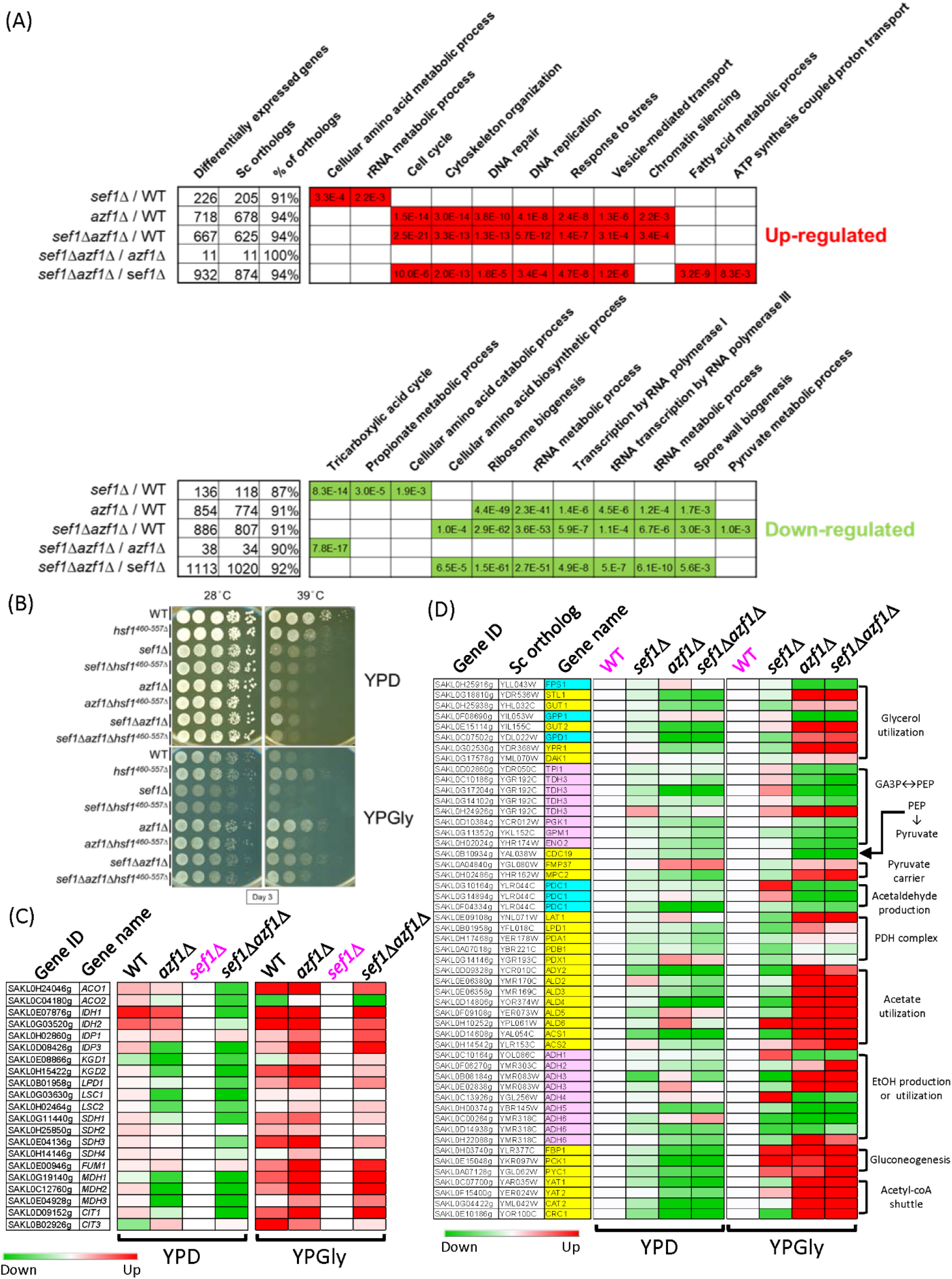
Phenotypic response to *azf1*Δ mutation in the YPGly condition. (A) Selected GO terms enriched under the YPGly condition. The adjusted *p*-values of the hypergeometric test with Bonferroni correction are shown. (B) The counteracting effect of hypomorphic *hsf1* on growth in response to *azf1*Δ mutation in both wild-type and *sef1*Δ backgrounds. *HSF1* is an essential gene in yeast (Solís et al. 2016), so a hypomorphic *hsf1* mutation with a truncated C-terminus (460-557 amino acids removed) (Sorger 1990) rather than a null mutant was used in *L. kluyveri* strains. (C) Partial compensation of defective TCA cycle gene expression in *sef1*Δ mutants by *azf1*Δ mutation under the YPGly condition. The heatmap was generated using the mean TPM ratio from RNA-seq data relative to the *sef1*Δ strain under each condition. (D) Transcriptional remodeling of the core glycerol utilization and auxiliary pathways in response to *azf1*Δ mutation under the YPGly condition. The heatmap was generated using the mean TPM ratio from RNA-seq data relative to the wild-type under each condition.

The “response to stress” GO term enriched in the *azf1*Δ-driven group encompasses 187 upregulated genes, including core heat-shock response genes and especially heat-shock proteins (HSPs) (Fig. S12A, see “Heat” and “Protein folding” sub-GO groups), implying a possible link between upregulated HSPs and “Gly-compensation” under heat-stressed conditions. HSPs play pivotal roles in protein folding and refolding, as well as in responses to heat and other stresses, and they are globally induced by the transcriptional regulator Hsf1, which is conserved across species (Veri et al. 2018). As expected, we found that a hypomorphic (diminished function) *hsf1* mutation partially abrogated the upregulation of representative HSP genes in response to *azf1*Δ (Fig. S12B), thereby partially limiting the “Gly-compensation” effect in both *azf1*Δ and *sef1*Δ*azf1*Δ mutant lines (Fig. 5B). That outcome indicates that the higher basal levels of HSPs induced by Hsf1 contribute to *azf1*Δ-induced heat resistance. Moreover, we identified a total of 212 genes related to ribosome/rRNA metabolism and a further 75 genes related to tRNA metabolism that were downregulated in response to *azf1*Δ (Fig. S13), demonstrating a typical yeast environmental stress response (ESR) in gene expression that involves coupling the upregulation of stress-defense genes with downregulation of ribosome biogenesis/protein synthesis genes (Gasch et al. 2000; Ho and Gasch 2015; Taymaz-Nikerel et al. 2016).

Furthermore, we noted that the TCA cycle GO term is not enriched among the downregulated genes of the *azf1*Δ-driven group, especially for *sef1*Δ*azf1*Δ/wild-type, unlike the *sef1*Δ-driven group (Fig. 5A). Indeed, deletion of *AZF1* not only partially restored expression of TCA cycle genes under the YPGly condition (Fig. 5C), but it also rescued cellular respiration [as assayed by triphenyl tetrazolium chloride (TTC) reduction assay] (Fig. S14A), thus reinforcing the “Gly-compensation” effect.

Apart from glycerol, both ethanol and acetate are common non-fermentable carbon sources preferred by yeast species during respiratory growth and that trigger a series of coordinated gene expression patterns (Turcotte et al. 2009). Interestingly, in contrast to the pronounced beneficial impact of glycerol, we found that acetate only provided a weak heat stress-specific benefit to the *azf1*Δ strains, whereas ethanol did not appear to exert any benefit (Fig. S14B). The same defective growth was observed in the post-diauxic growth phase of the *azf1*Δ mutant grown in YPD when ethanol was the major carbon source (Fig. S11B). To link this glycerol-specific trait to gene expression, we examined more closely the YPGly-based RNA profiles of the core non-fermentable carbon metabolic pathways of glycerol utilization (uptake and breakdown), pyruvate production, and pyruvate utilization (more specifically, the PDH-dependent respiratory pathway and the PDC-dependent fermentative pathway) (Fig. 5D). We present in Figure S14C the remodeled glycerol usage pathways and the predicted metabolic flux in *azf1*Δ cells (relative to wild-type) according to the differential gene expression summarized in Fig. 5D. In Figure S14D, we propose a model of glycerol-driven metabolic remodeling according to Fig. S14C. Our model is based on the notion that *azf1*Δ cells maintain a high intracellular concentration of glycerol and its derivatives due to increased uptake and restricted pyruvate production (from DHAP to pyruvate), with biased pyruvate flux fueling the TCA cycle due to strong competition for pyruvate between high-affinity mitochondrial pyruvate carriers plus the PDH complex and the low-affinity PDC complex (Pronk et al. 1996; Møller et al. 2004).

Hence, the loss-of-function *azf1* mutation induces “Gly-compensation” by: (1) increasing basal transcriptional levels of stress defense genes; (2) restoring TCA cycle gene expression; and (3) transcriptionally remodeling the metabolic pathways to favor glycerol utilization and more efficient respiration.

### *azf1*Δ cells display cell density-dependent “Dex-trade-off” traits and an evolutionary counteracting strategy

Intriguingly, during growth assays, we noticed unexpectedly that the “Dex-trade-off effect” in *azf1*Δ cells could be alleviated by gradually increasing the initial cell density (Fig. S15). More specifically, the maximal growth rate of *azf1*Δ cells under “Dex-trade-off” conditions was restored to a near wild-type level when growth was initiated at a higher inoculum size (Fig. 6A), hinting that a cell density-dependent mechanism plays a role in combatting the “Dex-trade-off effect”. To prevent extinction, *S. cerevisiae* cells may display cell density-dependent cooperative behavior to enable each other and their future generations to survive and replicate at high temperatures (Laman Trip and Youk 2020). To address if *L. kluyveri* cells can cooperatively help *azf1*Δ cells survive, we grew Azf1-sufficient (denoted *AZF1*) and *azf1*Δ strains independently in separate cultures or co-cultured together and measured the fitness of *azf1*Δ cells relative to the *AZF1* strain by counting viable cell numbers after 20-h of growth under the “Dex-trade-off” condition (Fig. S16A). Growth curves (Fig. S16B) revealed that the relative fitness of *azf1*Δ cells increased under co-culture conditions relative to the independent culture in both wild-type and *sef1*Δ backgrounds (Fig. 6B and 6C). Similar cooperative growth assays on an agar surface (Fig. S16C) generated results consistent with those from the equivalent aforementioned experiments in media (Fig. S16D and S16E). Taken together, these results indicate enhanced survival of *azf1*Δ cells under “Dex-trade-off” conditions can be facilitated by the presence of surrounding *AZF1* cells, mimicking the scenario by which low-frequency trade-off mutants can be maintained in a heterogeneous population.

**Figure 6.**
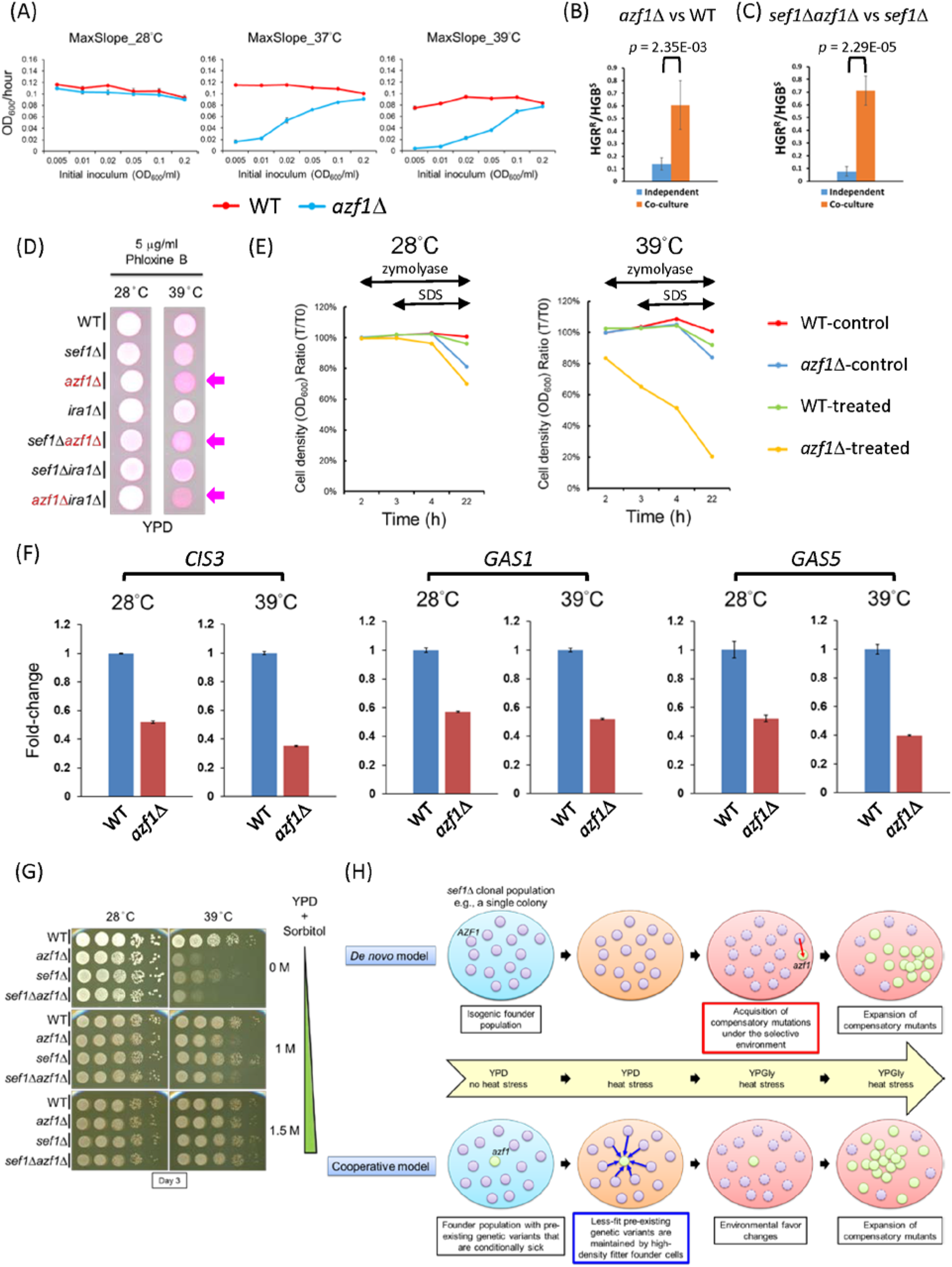
The trade-off phenotype of *azf1*Δ mutants is linked to cell wall integrity. (A) The effect of inoculum density on the mean maximal growth rate of the *azf1*Δ population in YPD under heat-stressed conditions. The growth curves were measured by using a Tecan plate reader with intermittent shaking. Error bars represent ± SD. (B) The *azf1*Δ cells are more persistent when co-grown with wild-type cells in liquid broth under the “Dex-trade-off” condition. (C) The *sef1*Δ*azf1*Δ cells are more persistent when co-grown with *sef1*Δ cells in liquid broth under the “Dex-trade-off” condition. For (B) and (C), statistical significance tests were carried out using unpaired Student’s t-tests. (D) Under the “Dex-trade-off” condition, *azf1*Δ cells were more severely stained by Phloxine B (pink color). Phloxine B is a negatively-charged dye that only stains cells intracellularly when cell membrane intactness has been compromised (Grosfeld et al. 2021). (E) The *azf1*Δ mutant is more sensitive to mild cell wall disruption. The cells pre-grown at indicated temperatures in YPD were first treated with 60 units/ml zymolyase at room temperature for 2 h, and then 0.1% SDS was added into the suspension to facilitate cell lysis. Cell intactness (y-axis) was calculated as the ratio of cell density (OD_600_/ml) at specific time-points relative to T_0_. Untreated samples were used as the control. (F) Expression of *CIS3* (SAKL0C05676g), *GAS1* (SAKL0H00550g), and *GAS5* (SAKL0F05456g) in response to *azf1*Δ mutation under the YPD condition. The relative fold-change of each gene is shown as 2^−ΔΔCT^, using *CDC34* (SAKL0D02530g) as the endogenous control and the ΔC_T_ value from the wild-type sample as the corresponding calibration value. Expression levels are displayed as means ± SD from three technical repeats. (G) The addition of sorbitol to the YPD medium to enhance osmolarity suppressed the “Dex-trade-off” phenotype of the *azf1*Δ mutants. (H) *De novo* and cooperative models can explain the rapid evolutionary repair of *sef1*Δ mutation-induced phenotypes by *AZF1* loss-of-function. In the *de novo* model, the *azf1* mutant newly forms under the selection brought about by *sef1*Δ-suppressive mutations. In the cooperative model, the *azf1* mutant pre-exists as a quasispecies in the founder population, its persistence being cooperatively supported by its fitter neighboring founder cells under the trade-off conditions, but it is then selected and undergoes population expansion to be dominant under favorable environmental conditions.

One mechanism by which cell density-dependent cooperative behavior operates in yeast is secretion of glutathione into the extracellular environment (Laman Trip and Youk 2020). The level of glutathione secretion required for growth under heat-stressed conditions depends on the cell population density before extinction. Although it remains unclear how extracellular glutathione helps cells to survive, one study has shown that a mild concentration of surfactants (such as sodium dodecyl sulfate, SDS) to increase cell permeability increased the extracellular concentration of glutathione (Wei et al. 2003). Other studies have indicated that glutathione is required for bacterial and yeast responses to osmotic stress (Smirnova et al. 2001; Jamnik et al. 2006) and that it serves as an osmotic driving force in bile formation (Jamnik et al. 2006). Taken together, these findings prompt the hypothesis that cell density-dependent cooperative behavior under heat stress may be linked to an osmotic imbalance of cells. Moreover, glutathione might be one but not the only factor contributing to this survival strategy.

Accordingly, we wondered if *azf1*Δ cells displaying cell density-dependent cooperative behavior also present an osmotic imbalance. Despite our gene expression data indicating that *azf1*Δ cells do not preferentially utilize ethanol, that finding is still insufficient to explain the ineffectiveness of ethanol (unlike glycerol) against heat stress. Consequently, we explored if ethanol is also a stressor of cells (Stanley et al. 2010), especially when cell wall integrity (CWI) is compromised (Udom et al. 2019). As expected, compared to wild-type, all *azf1*Δ cells not only displayed higher permeability to the dye phloxine B (Fig. 6D), but they were also more sensitive to mild cell wall disruption elicited by zymolyase treatment, especially when cells were pre-grown under heat-stressed conditions (Fig. 6E). Moreover, our finding that gene expression of a vital cell wall glycoprotein (*CIS3*) and twoβ-1,3-glucanosyltransferases (*GAS1* and *GAS5*) required for CWI in yeast (Tomishige et al. 2003; Mazáň et al. 2008) are all downregulated in *azf1*Δ cells (Fig. 6F) supports the notion that the *azf1*Δ mutants have a weakened CWI. Intuitively, a weakened cell wall (or outer membrane) will trigger hypo-osmotic stress in cells, leading to cell bursting if the condition is aggravated. Osmotic rebalancing by supplementing the medium with sorbitol can suppress growth defects caused by cell bursting (Lee et al. 1993; Serrano et al. 2006; Beese et al. 2009). We observed complete osmotic rebalancing in *azf1*Δ mutant cells upon adding 1.5 M of sorbitol into YPD (Fig. 6G), implying that the defective CWI of *azf1*Δ cells alters their optimal osmolarity required for growth and also contributes to their “Dex-trade-off” phenotypes. Interestingly, incorporating additional glutathione into the medium was unable to alleviate the growth defect of *azf1*Δ cells (data not shown), and we detected no clear differential gene expression related to glutathione in response to the *azf1*Δ mutation, indicating that further work is needed to identify the distinct mechanism by which Azf1 in *L. kluyveri* regulates this cell density-dependent cooperative behavior.

## Discussion

### Perturbing the Azf1 transcriptional network can rapidly compensate for disruption of the Sef1 transcriptional network

Our rapid suppressor development strategy allowed us to initiate an evolutionary repair experiment to determine responses to the *sef1*Δ mutation and to generate a large number of evolved lines carrying early and strongly beneficial mutations with the potential to compensate for the *sef1*Δ-caused defects. By performing selection on agar plates, clonal interference (Gerrish and Lenski 1998) among cells carrying beneficial mutations could be minimized, thereby circumventing a pervasive issue when evolving asexually-reproducing organisms in liquid broth (Barrick and Lenski 2013; Van den Bergh et al. 2018; McDonald 2019) and maintaining the early adaptive diversity of beneficial mutations (Levy et al. 2015; Blundell et al. 2019). Surprisingly, after phenotyping 240 suppressor clones from two independent lineages, we found that the majority (87.5%) displayed a consistent phenotypic pattern (“double-compensation” or “Dex-trade-off and Gly-compensation”). Sequencing three clones of each selected lineage and subsequent genetic verifications showed that they carried loss-of-function causal mutations of *IRA1* or *AZF1* (Fig. 2). That finding demonstrates clear evolutionary parallelism and unexpectedly restricted diversity arising from *sef1*Δ-suppressive mutations. We consider three possible reasons for that outcome. First, our suppressor colony picking strategy could have been biased because we picked the biggest colonies on the agar plates (Fig. S2) and clones of similar fitness most likely carry similar causal mutations, as described above for the beneficial *ira1*Δ mutation. Second, the specific type of driving forces may induce specific mutations, as supported by the triple beneficial effects of “restored respiration” (Fig. 5C and S14A), “elevated heat-shock defense” (Fig. S12), and “augmented glycerol utilization” (Fig. 5D) elicited by the *azf1* mutation, which together are required for cell survival under our heat-stressed YPGly condition. Moreover, given the potential for founder effects and epistasis (Barrick et al. 2010; Jerison et al. 2017; Wünsche et al. 2017; Rojas Echenique et al. 2019), we cannot exclude the possibility that the *sef1*Δ founder genotype constrained or shaped the fitness landscape. Third, pre-existing low-frequency beneficial mutations in the founder population may have been selected, with this latter scenario being supported by our identification of the presence of identical mutation types in separate suppressor clones. Indeed, we identified four such cases (Fig. 2C and 2D), including the Val2130_Val2137Δ and Arg39Cys mutations of *IRA1* between 28°C-Evo clones, as well as the Asp485Tyr and Cys545Phe mutations of *AZF1* between 39°C-Evo clones. Identical mutations are unlikely to form independently by chance in an asexual population, so we conclude that these mutations may have pre-existed in the quasispecies founder population (Fig. S1E), acting as rapid adaptive solutions to the changing environment (Teng et al. 2013).

### Glucose, Ras1-PKA signaling, and Azf1

The beneficial effects of the *ira1* loss-of-function mutation arise from hyperactive Ras-PKA signaling, leading to uncontrolled cell growth in the absence of the yeast growth-promoting factor glucose, reminiscent of tumor cell growth (Cazzanelli et al. 2018). That mutation exerted a generalist impact, eliciting a “double-compensation” effect irrespective of the conditions used in our experiments. In contrast, the *azf1* loss-of-function mutation is pleiotropic in terms of the aforementioned triple beneficial effects and its deleterious impacts relating to mildly defective core carbohydrate metabolism (Fig. 4 and S8), amino acid metabolism (Fig. S11), and CWI (Fig. 6). All those defects only arose under the YPD condition and they were exacerbated by heat stress, indicating that they represent life history “trade-offs” (Garland 2014) and that the *azf1* mutant cells are typical specialists (Van den Bergh et al. 2018).

Interestingly, although Azf1 appears functional under both glucose and glycerol conditions, we believe that glucose is the environmental factor sensed by Azf1, i.e., Azf1 regulates gene expression in response to the presence or absence of glucose. This speculation is supported by the facts that: (1) Azf1 activity is regulated by the Ras1-PKA pathway (Fig. 3), which is mainly glucose-responsive in yeast (Conrad et al. 2014; Cazzanelli et al. 2018); (2) *azf1*Δ triggered the opposite expression pattern between stress response and ribosomal genes (Fig. S12A and S13). This opposite expression pattern is also regulated by the PKA pathway in response to the glucose availability (Conrad et al. 2014); and (3) the presence of glycerol in YPD did not convert “Dex-trade-off” to “Gly-compensation” (Fig. S17), implying that glucose is the trigger. Elucidating the molecular components involved in these processes is an interesting topic for future exploration.

### Cooperation maintains trade-off genotypes

We have revealed that pleiotropic defects (misregulation of multiple TCA genes due to the *sef1*Δ mutation in this study) can be simultaneously compensated in a short timeframe simply by means of another pleiotropic change (i.e., *AZF1* loss-of-function). Despite potentially being maintained by balancing selection (Mérot et al. 2020), this new pleiotropic change may remain concealed by accompanying trade-offs within a short evolutionary timescale, especially in unstable environments (Chen and Zhang 2020). Here, we have demonstrated cell density-dependent mitigation of such trade-offs, whereby *azf1*Δ individuals can persist when densely surrounded by founder cells or until the density of *azf1*Δ progeny exceeds a threshold (Fig. 6A, 6B, 6C, S16D, and S16E). Although the underlying mechanism remains unknown, this “cooperative persistence” enhances the adaptive potential not only of *de novo azf1* mutants when unfavorable environments are encountered but also of pre-existing *azf1* mutants before experiencing selection (Fig. 6H), enabling a contingent secondary adaptive step to repair the trade-offs (Aggeli et al. 2021).

Frequency-dependent selection (FDS) describes how the fitness of a genotype or phenotype in a population is related to its frequency in the population (Ayala and Campbell 1974). Positive FDS is a selection regime whereby the fitness of a phenotype increases with its frequency. The cell density-dependent mitigation of trade-offs in *azf1*Δ cells represents a simple case of positive FDS. In early models, positive FDS was predicted to reduce diversity due to systematic loss of rare polymorphisms. However, more recent theories predict that positive FDS maintains rather than reduces diversity when populations patchily occupy spatially structured habitats (Molofsky et al. 2001; Molofsky and Bever 2002; Rendueles et al. 2015). Our evolutionary repair experiments, with characteristics of changing environments (fermentative vs respiratory; normal vs heat-stressed), local habitats (growth on agar plates), and cell density-dependent cooperation, is an intuitive example showing that positive FDS maintains microbial diversity and facilitates evolution.

The *sef1*Δ−suppressive mutations of *AZF1* compensate for perturbed expression of vital TCA cycle genes under selective conditions, reflecting a type of evolutionary tinkering and stabilizing selection (Lavoie et al. 2010; Signor and Nuzhdin 2018; Signor and Nuzhdin 2019). The multiple beneficial and trade-off mutations that accumulate as a consequence of ongoing evolutionary repair processes represent a good source of molecular diversity (LaBar et al. 2020). From the perspective of the macroevolution of transcriptional networks, our study illuminates the process of incipient transcriptional rewiring and formation of complex traits during evolution of transcriptional networks.

## Materials and Methods

### Genome resources

The genome sequence of *Lachancea kluyveri* (CBS 3082) was originally from Bioproject accession number PRJNA1445. The sequence and improved annotation can be downloaded from the GRYC (Genome Resources for Yeast Chromosomes; http://gryc.inra.fr) database (Brion et al. 2015; Hsu et al. 2021). The sequences and structures of the *L. kluyveri* MATa and MATα loci were as per contigs under accession numbers AACE03000005 (Consortium et al. 2009) and AACE01000014 (Butler et al. 2004), respectively.

### Strains and plasmid construction

To avoid confusion, we have adopted the term “wild-type” to describe the parental genotypes used in our study, each of which may carry auxotrophic mutations or specific genetic manipulations, and they acted as reference strains for all comparative experiments unless otherwise stated. The *L. kluyveri* strains, primers, and plasmids used in this study are listed in Table S13. All DNA fragments used in cloning were PCR-amplified using Phusion High-Fidelity DNA polymerase (F530L, Thermo Fisher Scientific, USA). The PCR products and restriction digested products were purified by PCR cleanup or gel extraction using an AccuPrep^®^ PCR/Gel Purification Kit (K-3037, Bioneer, Korea). DNA cloning into plasmids was achieved by using ligation (T4 DNA ligase, M180A, Promega, USA) or by means of an In-Fusion^®^ HD Cloning Kit (639650, ClonTech by Takara, Japan). All plasmids were extracted and purified using a Presto^TM^ Mini Plasmid Kit (PDH300, Geneaid, Taiwan). Successful strain constructions were confirmed by genomic DNA extraction, as described previously (Hsu et al. 2011), followed by PCR diagnostics using a homemade Taq DNA polymerase for genotyping. For gene deletions, the DNA fragments of each deletion module consisting of a selection marker flanked by 5’ and 3’ sequences homologous to the target locus were created by overlap-extension-PCR (Shevchuk et al. 2004). The genes deleted in this study are: *SEF1* (SAKL0F12342g), *CHS3* (SAKL0C11726g), *AZF1* (SAKL0E10714g), *IRA1* (SAKL0C10340g), *PDE1* (SAKL0F00638g), *PDE2* (SAKL0D14344g), and *BCY1* (SAKL0F07788g).

For lexA and TAP tagging of Azf1, we used a modified (Hsu et al. 2021) SAT1 flipper method (Reuß et al. 2004). The TAP tagging vector pSFS2A-TAPSacISacIIMut-28 has a second SacII site near the C-terminus of the TAP sequence, rendering cloning at the 3’ multiple cloning site (MCS) problematic. To create a new TAP tagging vector, we mutated the second SacII site by amplifying it from pSFS2A-TAPSacISacIIMut-28 using the primer pair TAP-1-ApaI and TAP-2-SacIIMut-XhoI, and then ligating the product between ApaI-XhoI sites in the pSFS2A vector to generate pSFS2A-TAP-New-2. The 978-bp C-terminal sequence (+1480 to +2457 of the ORF without the STOP codon) of *AZF1* was cloned into the pSFS2A-lexA-5 or pSFS2A-TAP-New-2 vectors between the KpnI-ApaI sites using In-Fusion^®^, and the 411-bp 3’-UTR (terminator, +1 to +411 from the STOP codon) sequence of *AZF1* was ligated between the SacII-SacI sites to generate the lexA- or TAP-tagging plasmid for *AZF1*. The KpnI-SacI-digested fragments from both tagging plasmids were used to transform cells. The SklexAOPlacZ-1 reporter strain and derivatives were transformed to create chromosome-integrated Azf1 one-hybrid strains, whereas the JYL1897 wild-type strain and derivatives were transformed to generate Azf1-TAP strains. To produce plasmid-based Azf1 one-hybrid strains, the entire 2460-bp *AZF1* ORF with STOP codon was cloned in-frame into pRS41H-lexA-3 between the BamHI-NotI sites downstream of the lexA sequence, thereby generating pRS41H-lexA-SkAzf1-1, and the plasmid was transformed into the SklexAOPlacZ-1 strain and derivatives. For Sef1, previously described one-hybrid and TAP-tagged strains were used (Hsu et al. 2021).

To create the *RAS1^G20V^* hyperactive one-hybrid strains, we used the SAT1 flipper method. Gly20 of *RAS1* (SAKL0E10252g) was first mutated to Val by overlap-extension-PCR-based site-specific mutagenesis (Sambrook 2001). The 1395-bp 5’-flanking region of the *RAS1^G20V^* fragment (−424 from ATG to STOP) and its 297-bp 3’ flanking region (+89 to +385 from STOP codon) were then ligated into the 5’ and 3’ MCSs of the pSFS2A vector between the ApaI-XhoI sites and the SacII-SacI sites, respectively. To transform cells, the ApaI-SacI-digested fragments from the pSFS2A-Ras1G20V-14 plasmid were used during transformation to replace the wild-type allele.

To create the *hsf1* (SAKL0A04576g) hypomorphic strains, we used the DAmP (Decreased Abundance by mRNA Perturbation) (Breslow et al. 2008) and decrease-of-function *hsf1* mutant (Sorger 1990). In brief, the 475-bp DNA fragment of C-terminal-deleted (Δ460-557 aa) *hsf1* (SAKL0A04576g, +903 to +1377 from ATG), a drug marker (KanMX6 or NatMX4), and the 355-bp 3’ flanking region (+27 to +381 from the STOP codon) were PCR-fused and used during transformation to replace the wild-type allele.

For mating-type switching from MATa to MATα, we again used the SAT1 flipper method. The 499-bp 5’ flanking region of the MATα locus near the *DIC1* locus, together with the 3206-bp fragment composed of the 2587-bp MATα locus and the 619-bp 3’ flanking regions of the *SLA2* locus, were ligated into pSFS2A vector between the KpnI-ApaI sites and NotI-SacII sites, respectively, to create the mating-type switching plasmid pSFS2A-5f-SkMATα-1. To transform cells, the KpnI-SacII-digested fragments from the pSFS2A-5f-SkMATα-1 plasmid were used.

All yeast transformations were performed by using electroporation and transformed cells were then selected as described previously (Hsu et al. 2021). Notably, since the *Candida albicans MAL2* promoter on the SAT1-FLIP cassette is leaky in *L. kluyveri*, the *L. kluyveri* SAT1-FLIP transformants could only be selected and propagated in YPD+Nou10 (10 μg/ml Nouseothricin) plates during strain construction procedures. The integrated SAT1-FLIP cassette does not support efficient growth of *L. kluyveri cells* in liquid broth with Nouseothricin.

### Media, important chemicals, and growth conditions

All media and important chemicals used in this study are listed with abbreviations in Table S13. All agar plates contained 2% agar. The basal YP (1% yeast extract and 2% peptone)-based media were sterilized by autoclaving, except for YPEtOH (YP + 2% ethanol). YPEtOH and all synthetic media were sterilized by filtering through 0.2 μm cup filters (Nalgene™ Rapid-Flow™ Sterile Single-Use Bottle Top Filters, 595-4520; Thermo Fisher Scientific, USA). Some modifications to the recipes for synthetic media are noted in Table S13. All cultures in culture tubes were grown on a drum roller rotating at 65–80 rpm. The normal growth temperature was 28–30°C. Heat stress was applied at 37−39°C, as indicated in the experiments.

### Evolutionary repair experiments by suppressor development

The entire procedure is summarized in Fig. S2. In detail, the MATa (SkSef1KChs3HA2) and MATα (SkSef1KChs3Hα1) *SEF1*-deleted founder strains (*sef1*Δ::KanMX6, *chs3*Δ::HphMX4, *ura3*^−^) were inoculated into a 20-ml YPD culture and grown at 30°C. After overnight growth, the cells were harvested and resuspended in 2 ml sterile ddH_2_O at a cell density of ∼40 OD_600_/ml. Approximately 100−120 μl of cell suspensions containing 5 OD_600_ cells (∼10^8^ CFU) were plated onto fresh YPGly plates, with 3−5 repeats per condition. The plates were incubated at 28, 37, 38, or 39°C. After 8 days, outcompeting colonies were re-streaked on YPGly plates and selected for another 5 days at 28, 37, 38, or 39°C, respectively. Only one large single colony was picked from each re-streaked population, and then the clones were streaked on YPD+HGB plates and incubated at 28°C for 2−3 days to confirm the original *chs3*Δ::hphMX4 genotype and to exclude contamination. The HGB-resistant clones were picked and inoculated into 1-ml YPD+G418/well cultures in U-shaped 96-deep-well plates (Nunc™ 96-Well Polypropylene DeepWell™ Storage Plates, 278743, Thermo Fisher Scientific) covered with sterile breathable sealing film (BF-400-S, Axygen by Coring, USA). The deep-well plates were incubated with shaking (Micromixer MX4, FINEPCR, Korea) at 30°C for 24 h to amplify the cells, before confirming the original *sef1*Δ::KanMX6 genotype. The cells from saturated cultures were spun down and concentrated to ∼130 μl, before adding 50 μl of 85% sterile glycerol and transferring them to 96-well plates (Tissue Culture Testplate 96 wells F-bottom, 92096, TPP, Switzerland) sealed with adhesive foil (UTI 3101, Smartgene by Ultra Violet, Taiwan) for −80°C storage.

### Genetic analyses by tetrad dissection

To mate yeast cells, 750 μl (∼4 OD_600_ containing ∼1×10^8^ CFU) each of MATa and MATα cells from YPD overnight cultures grown at 28°C for at least 20 h were mixed and harvested. The mixed cell pellets were washed once with 1 ml sterile ddH_2_O, resuspended in 10 μl sterile ddH_2_O, and then spotted onto dry YPD plates. After growing at 28°C for 5-7 h, the cells from the spots were separated by streaking, and candidate zygotes were picked by using a Tetrad Dissection System [Nikon Eclipse 50i Microscope (Nikon, Japan) equipped with a TDM50^TM^ Micromanipulator (Micro Video Instruments, Inc., USA) and a dissection needle (NDL-010, Singer Instruments, UK)]. The candidate zygotes were incubated on YPD plates at 28°C for 2-3 days and the heterozygous genotypes at the MAT loci were confirmed by gDNA extraction followed by PCR diagnosis for both MATa and MATα alleles.

For sporulation, 1 ml zygotes from YPD overnight culture were pelleted down, washed once with 1 ml sterile ddH_2_O, and then inoculated into 3 ml Spo medium (2% potassium acetate pH 9, optional: +1% adenine and 1% tryptophan) at 23−25°C. After 5-7 days, the asci from 50 μl sporulation culture were harvested, resuspended in 10 μl of 1 mg/ml zymolyase working solution diluted from 40 mg/ml zymolyase stock solution [Zymolyase^®^ 100T (07665-55, Nacalai Tesque Inc., Japan), 1X TE (10 mM Tris-Cl, 1 mM Na2EDTA pH 8), 5% glucose] in sterile 1 M sorbitol or ddH_2_O, and digested at room temperature for 5−8 min. The digested asci were streaked on dry YPD plates and the four spores from each tetrad were picked and dissected onto the space of the same YPD plate. The dissected sister spore sets were incubated at 28°C for 2-3 days for further genotyping and phenotyping.

Notably, all strains used for tetrad dissection were in the *chs3*Δ background, which breaks the interspore bridges between sister spores in asci (Coluccio and Neiman 2004) and allows separation of sister spores, making *L. kluyveri* tetrad dissection achievable (Sigwalt et al. 2016).

To sequence the *AZF1* locus, the entire *AZF1* ORF (−21 from ATG to +17 from STOP) was PCR-amplified using Phusion High-Fidelity DNA polymerase (F530L, Thermo Fisher Scientific) with the primer pair SkAzf1-UpATG-1 and SkAzf1-STOPdown-2 and an annealing temperature of 52°C. The product was purified by PCR cleanup using an AccuPrep^®^ PCR/Gel Purification Kit (K-3037, Bioneer), before being subjected to Sanger sequencing with primer SkAzf1-555-1-seq 1 or SkAzf1-1151-1-seq 1. To sequence the *IRA1* locus, the second half of the *IRA1* ORF (+4744 from ATG to +40 from STOP) was PCR-amplified as detailed above, but with the primer pair SkIra1-4744-1-seq 1 and SkIra1-STOPdown-2 and an annealing temperature of 58°C, before undergoing purification and Sanger sequencing with primer SkIra1-5356-1-seq 1 or SkIra1-5946-1-seq 1. All primers used are listed in Table S13.

### Phenotypic assays

The spot assays, growth curve assays, and TTC reduction assays were performed as described previously (Hsu et al. 2021) with minor modifications. For the spot assays, cells grown in YPD overnight at 28°C were harvested by centrifugation (Eppendorf 5810R centrifuge; A-4-62 rotor; 1500 *g*; 3 min; 25°C) and serially diluted to the desired cell density with sterile ddH_2_O. Each dilution (10^7^–10^3^ CFU/ml) was spotted onto agar plates (5 μl/spot, about 10^5^–10^1^ CFU/spot) and incubated at 28°C or other temperatures as indicated. The plates were scanned using a scanner (Epson Perfection V800 Photo) and images were recorded after 3 days or as otherwise indicated.

For the growth curve assays, cells grown overnight in YPD (or other media as indicated) at 28°C were inoculated into 120 μl YPD medium in a 96-well plate (Tissue Culture Testplate 96 wells F-bottom, 92096, TPP) at a cell density of 0.2 OD_600_/ml at indicated temperatures. Cell growth was measured at OD_595_ in “2×2 multiple reads per well” mode every 12 min using a Tecan plate reader (Infinite 200 PRO, Tecan, Switzerland). Tecan software Magellan Version 7.2 was used for data acquisition and analyses. For the Magellan method, we selected the plate definition “[TPP96ft]-Tecan Plastic Products AG6 Flat Transparent”. Each 12-min cycle comprised 3 min reading, 1 min shaking, 3 min standing, 1 min shaking, 3 min standing, and 1 min shaking. For the glycolysis-inhibited medium, 0.1% 2-deoxy-D-glucose (D8375, Sigma) was added to the YPD medium. For amino acid pre-starvation, we used SM+2X Ura medium (SM supplemented with 80 mg/l uracil) given that we employed *ura*^−^ strains in this study. Pre-starvation was performed by inoculating a single colony from a YPD plate into a 5-ml SM+2X Ura culture and allowing it to grow for 23 h at 28°C. For cell density-dependent growth assays, growth curves using different initial cell densities (0.005, 0.01, 0.02, 0.05, 0.1 or 0.2 OD_600_/ml) in YPD were measured for 24 h at 28, 37, or 39°C. Maximal growth rates were determined by calculating the maximum slope of each growth curve based on 30−40 time-points in Magellan software, as illustrated in Fig. S1C.

For the TTC reduction assays, cells grown overnight in YPD at different temperatures as indicated were harvested and diluted to a cell density of 1 OD_600_/ml in sterile ddH_2_O. Then, 5 μl of each cell suspension (∼10^5^ CFU) was spotted on YPGly plates. The plates were incubated at 28°C for 2 days, before being overlaid with 20 ml melted TTC agar (0.1% TTC, 2% agar in sterile PBS [8 g/l NaCl, 0.2 g/l KCl, 1.44 g/l Na_2_HPO_4_, 0.24 g/l KH_2_PO_4_ pH 7.4]) and allowed to solidify for 20 min at room temperature. The plates were incubated at 28°C for red color development. The plates were scanned after 2.5, 8, and 24 h of color development using a scanner (Epson Perfection V800 Photo).

For the Phloxine B permeability assays, cells grown overnight in YPD at 28°C were harvested by centrifugation (Eppendorf 5810R centrifuge; A-4-62 rotor; 1500 *g*; 3 min; 25°C) and diluted to a cell density of 1 OD_600_/ml in sterile ddH_2_O. Then, 5 μl of each cell suspension (∼10^5^ CFU) was spotted on YPD plates containing 5 μg/ml Phloxine B (P4030, Sigma). The plates were incubated at 28 or 39°C and scanned after 3 days of pink color development using a scanner (Epson Perfection V800 Photo). To intensify differences in color, the images were processed uniformly by increasing contrast and brightness by 40%.

For the cell wall disruption sensitivity assays, cells grown in YPD for 16 h at 28°C were harvested and re-inoculated into 5 ml of YPD at a cell density of 0.2 OD_600_/ml for 24 h at 28 or 39°C. The cells were harvested by centrifugation (Eppendorf 5810R centrifuge; A-4-62 rotor; 1500 *g*; 3 min; 25°C) and diluted at cell densities of 0.6 OD_600_/ml with 1 ml of pH 7.6 Tris-Cl buffer containing 60 units/ml Zymolyase^®^ 100T (07665-55, Nacalai Tesque Inc.) in a disposable 2-ml photometer cuvette (112117, LP Italiana SPA, Italy). The cell walls were digested initially for 2 h in the Zymolase-hosting buffer, before further digestion using 0.1 % SDS at room temperature, with gentle pipetting every 30 min. OD_600_ at 0, 2, 4, and 22 h was measured using a Beckman DU^®^ 800 UV/Visible spectrophotometer (Beckman Coulter, USA) to evaluate intact cells. Cells with disrupted cell walls lyse in the presence of SDS due to hypoosmolarity, reducing the OD_600_ value in the cell suspension. The ratio of OD_600_ at each time-point relative to time zero (0 h) was calculated to represent sensitivity to cell wall disruption, with a lower ratio reflecting higher sensitivity.

For the cooperative growth assays in liquid broths (Fig. S16A), cells grown overnight in YPD (20 h) at 28°C were harvested and diluted to a cell density of 1 OD_600_/ml in sterile ddH_2_O. For the “independent cultures”, *AZF1* (reference strain, HGB-sensitive) and *azf1*Δ (test strain, HGB-resistant) cell suspensions were diluted independently to a cell density of 0.01 and 0.05 OD_600_/ml, respectively, in a 1-ml YPD culture. For the “co-culture”, the *AZF1* and *azf1*Δ cell suspensions were mixed at cell densities of 0.01 and 0.05 OD_600_/ml, respectively, in the same 1-ml YPD culture. Cell growth was measured at 39°C using a Tecan plate reader (Infinite 200 PRO, Tecan), as described above. After 20 h of growth, 50 μl of each of the “independent cultures” was mixed and resuspended in 900 μl sterile ddH_2_O, and then 100 μl of the “co-culture” was resuspended in 900 μl sterile ddH_2_O. Two suspensions (∼0.4 OD_600_/ml) were serially diluted and plated onto YPD plates at an estimated density of ∼10^2^ CFU/plate. The YPD plates were incubated at 28°C for 2 days, before being replicated onto YPD+HGB plates and incubated for 1 day. The ratio of *azf1*Δ (CFU^HGB-resistant^) to *AZF1* (total CFU^YPD^ – CFU^HGB-resistant^) cells was calculated from six technical repeats.

For the cooperative growth assays on agar plates (Fig. 16C), cells grown overnight in YPD (20 h) at 28°C were harvested and diluted in sterile ddH_2_O at cell densities of 1 OD_600_/ml for the wild-type and 5 OD_600_/ml for the *azf1*Δ mutant. For the “independent cultures”, 5 μl of the wild-type (reference strain, HGB-sensitive, ∼10^5^ CFU) and *azf1*Δ (test strain, HGB-resistant, ∼5×10^5^ CFU) cell suspensions were spotted independently onto a YPD plate. For the “co-culture”, 5 μl of the mixed wild-type and *azf1*Δ cell suspension containing the same CFU as described above was spotted onto the same YPD plate. After incubation at 39°C for 3 days, 7×7 mm areas of gel hosting cell colonies were sliced out and rehydrated with 1 ml sterile ddH_2_O at room temperature. After 1 h, the cells were washed down five times by means of mild pulse vortexing. Suspensions were transferred to new 2-ml tubes and the wash step was repeated. The new suspensions were combined with the previous one. Two slices from the “independent cultures” were rehydrated and resuspended together, whereas only one slice from the “co-culture” was processed. Two suspensions (∼3 OD_600_/ml for the “independent culture” and ∼1.6 OD_600_/ml for the “co-culture”) were then serially diluted and drop-plated. Each dilution (10^6^–10^3^ CFU/ml) was spotted onto the YPD and YPD+HGB plates (5 μl/spot, ∼10^4^–10^1^ CFU/spot) and incubated at 28°C for 2−3 days. Spots with <50 CFU were counted. Notably, instead of using the HGB-resistant to HGB-sensitive ratio, we used the frequency of *azf1*Δ cells (CFU^HGB-resistant^/total CFU^YPD^) to represent the level of cooperative growth, thereby reducing technical variations caused by the considerable difference in viability between *AZF1* and *azf1*Δ cells on agar plates. The frequencies were calculated from six technical repeats. Due to their lower fitness at 39°C, for cases in the *sef1*Δ background (*sef1*Δ, HGB-sensitive; *sef1*Δ*azf1*Δ, HGB-resistant), the cell densities of initial cell suspensions were modified to 2 OD_600_/ml for the *sef1*Δ mutant and 4 OD_600_/ml for the *sef1*Δ*azf1*Δ mutant.

### Beta-galactosidase assays

LacZ expression levels for the one-hybrid assays were quantified using liquid β-galactosidase assays, as described previously but with some modifications (Hsu et al. 2021). In brief, early log-phase (0.5–1.0 OD_600_/ml) cells from subcultures of indicated media were assayed. All subcultures were initiated with 0.2 OD_600_/ml of inoculum from overnight-grown cells. One-hybrid strains from the chromosome-integrated system were subcultured for 4.5 h, whereas the strains from the plasmid-based system were subcultured for 5.5 h in the presence of 200 μg/ml HGB. Due to the relatively weak activation of Azf1, more cells (∼1 OD_600_ cells) per reaction and a longer incubation time (90 min) at 37°C were required for colorimetric reactions. LacZ levels are displayed as Miller units.

### Genomic DNA extraction for whole-genome sequencing

Pure genomic DNA was extracted via the phenol-chloroform method, followed by RNase treatment, as described previously but with modifications (Hsu et al. 2011). In brief, ∼100 OD_600_ cells (from a 20-ml YPD overnight culture) per sample were extracted. The cells were resuspended in 200 μl gDNA lysis buffer [2% Triton X-100, 1% SDS, 100 mM NaCl, 10 mM Tris-Cl (pH 8), 1 mM Na_2_EDTA] and transferred into a 2-ml breaking tube (72.693, Sarstedt) containing 250 μl PCIA (phenol:chloroform:isoamyl alcohol=25:24:1, 0.1% 8-quinolinol, pH 8) and a 500-μl volume of glass beads (11079105, BioSpec Products). The cells were lysed by using a FastPrep-24™ 5G Homogenizer (MP Biomedicals™) at a speed of 6 m/s for 40 sec. The aqueous phases were then extracted again using 200 μl PCIA (pH 8). RNase treatment involved adding 10 μl of 10 mg/ml RNase A (R5503, Sigma) into a 400-μl nucleic acid suspension in TE buffer (10 mM Tris-Cl, 1 mM Na_2_EDTA, pH 8) and incubating at 37°C for 1 h. The crude gDNA was then precipitated by adding 1 ml of 99% ethanol with 20 μl of 3M sodium acetate (pH 5.2), followed by mild mixing for 30 min. The nucleic acid pellets were rehydrated in 200 μl ddH_2_O at 65°C for 10 min and then purified again by using a Zymo Genomic DNA clean & concentration^TM^ kit-25 (Zymo Research, USA) to remove the small DNA fragments. All procedures followed the manufacturer’s instructions, but with the modification of a two-cycle wash step using 500 μl wash buffer. The final gDNA was eluted with 50 μl of 65°C elution buffer. DNA concentrations were measured using a Qubit™ dsDNA BR Assay Kit (Q32853, Invitrogen by Thermo Fisher Scientific). The quality of the gDNA was checked using the 2200 TapeStation system (G2964AA), Genomic DNA ScreenTape (5067-5365), and Genomic DNA Reagents (5067-5366) obtained from Agilent Technologies.

### DNA-seq analyses

Isolated genomic DNAs were fragmented using a Covaris M220 Focused-ultrasonicator (Covaris, USA), and DNA libraries were prepared using a Truseq DNA PCR-free HT kit (Illumina, USA). Two founders and 12 suppressor clones were sequenced to an estimated average depth of 50-fold coverage by paired-end 150 bp read length following the protocol of the NextSeq 500 Mid output v2 (300 cycles) sequencing kit on an Illumina NextSeq 500 System controlled by the NextSeq Control Software (NCS v2.2.0, Illumina). Real-Time Analysis (RTA v2.4.11, Illumina) software was used to generate the BCL files containing base calls and associated quality scores (Q-scores). The bcl2fastq Conversion Software (v2.17, Illumina) was used to demultiplex sequencing data and convert the BCL files into FASTQ files, which were then processed by FastQC (www.bioinformatics.babraham.ac.uk/projects/fastqc) and summarized in MultiQC (Ewels et al. 2016) to generate a QC report for comparison.

Raw reads were mapped to the reference CBS 3082 genome using the Burrows-Wheeler Aligner (BWA, v0.7.17) and the BWA-MEM algorithm (Li and Durbin 2009; Li 2013) with default parameters to obtain the BAM files. Mapping rates and sequencing coverages were determined in Samtools v1.9 (Li et al. 2009). Variant calling was performed on the BAM files using GATK v4.1.2.0 (McKenna et al. 2010). According to “hard-filtering” recommendations for germline short variant (SNP and INDEL) discovery described in the GATK website (https://gatk.broadinstitute.org/hc/en-us/), we applied the following filtering settings with some modifications for SNP calling: “QD < 2.0 || FS > 60.0 || MQ < 40.0 || MQRankSum < −12.5 || ReadPosRankSum < −8.0 || SOR > 4.0”, and for INDEL calling: “QD < 2.0 || FS > 200.0 || ReadPosRankSum < −20.0”, to obtain VCF files. Only variants existing in the suppressor clones but not in their founder strain were retained for subsequent analyses. The suppressor-unique variants were then filtered according to sequencing depth (DP) >20 and allele frequency (AF) ≥0.5 to get the final variant spectra. SnpEff v4.3t (Cingolani et al. 2012) was used to annotate and predict the effects of suppressor-unique variants on genes. The final unique variants with annotations are listed in Table S4.

### RNA extraction and DNase treatment

Approximately 10 OD_600_ of early log-phase (0.5–1.0 OD_600_/ml) cells from subcultures grown from 0.2 OD_600_/ml of inoculum for 5.5 h under the indicated conditions were harvested. Total RNA was extracted by means of the phenol-chloroform method and treated with the TURBO DNA-freeTM kit (AM1907, Ambion, Invitrogen by Thermo Fisher Scientific), as described (Hsu et al. 2021) but with small modifications. In brief, the cells were resuspended in 500 μl nuclease-free and ice-cold RNA lysis buffer and then transferred into a 2-ml breaking tube (72.693, Sarstedt, Germany) containing 250 μl ice-cold acidic PCIA and a 500-μl volume of glass beads (11079105, BioSpec Products, USA). The cells were lysed using a FastPrep-24™ 5G Homogenizer (MP Biomedicals™, USA) at a speed of 6 m/s for 40 sec at room temperature. The aqueous phases were then extracted by two rounds of 200-μl ice-cold acidic PCIA treatment, and the nucleic acids were obtained after 2 h of ethanol precipitation. RNA quality was checked using a Bioanalyzer 2100 instrument (Agilent Technologies, USA) with an RNA 6000 Nano LabChip kit (Agilent Technologies). The DNase-treated RNAs were stored at 80°C for further RT-qPCR or RNA-seq analyses.

### Reverse transcription (RT) and quantitative PCR (qPCR) analyses

RT-qPCR was performed as described previously but with some modifications (Hsu et al. 2021). In brief, cDNA synthesis was conducted in the presence of different rRNasin^®^ Ribonuclease Inhibitors (N2511, Promega, USA) at a concentration of 1 μl per 20-μl reaction. qPCR was performed using the model QuantStudioTM 12K Flex Real-Time PCR System (Applied Biosystems by Thermo Fisher Scientific). Primers used for qPCR are listed in Table S13. *L. kluyveri CDC34* transcripts (SAKL0D02530g) were used as an endogenous control for qPCR. The average ΔΔC_T_ and standard deviation were determined from at least three technical repeats. The relative fold-change of each gene is shown according to the 2^−ΔΔCT^ method.

### RNA-seq analyses

For each strain and condition, total RNA was extracted from three biological replicates. RNA-seq libraries were prepared using a TruSeq Stranded mRNA HT Sample Prep Kit (Illumina) on 2 μg total RNA and sequenced by single-end 75 bp read length following the protocol of the NextSeq 500 High Output kit v2.5 (75 cycles) sequencing kit on an Illumina NextSeq 500 System. Total read numbers were estimated to be 20 million reads per sample. Raw reads were quality-trimmed using Trimmomatic v0.38 (Bolger et al. 2014) with options: “ILLUMINACLIP:2:30:10 LEADING:3 TRAILING:3 SLIDINGWINDOW:4:15 MINLEN:36”. The relative abundance of each gene in units of Transcripts Per Million (TPM) for each sample was quantified using Salmon v1.4.0 (Patro et al. 2017) with options: “--numBootstraps 200 -- validateMappings --gcBias”. Tximport (Soneson et al. 2015) by default was used to convert the estimated transcript abundance files from Salmon to a DESeq2-compatible dataset. DESeq2 (Love et al. 2014) was used for the analysis of differential gene expression in R v4.0.4 with default settings. In DESeq2, the *p*-values attained by the Wald test were corrected for multiple testing using the Benjamini and Hochberg method. Genes with mean TPM < 1 under all conditions were omitted from our analyses. Differences in gene expression with adjusted *p*-values <0.05 and fold-change of at least 2 were considered significantly different. All results in the standard DESeq2 output format and the TPM of each gene from three technical repeats are presented in Table S1, S2, and S5-S12.

The heat maps of selected gene expressions were created using the mean TPM ratio relative to a reference sample, as indicated. The green-white-red color scale ranging from the 5^th^ to 90^th^ percentiles of the selected values was generated in Excel 2016 using the conditional formatting function. The while color represents the median with a value of ∼1, indicating no differential expression.

### Gene ontology (GO) enrichment analyses and manual simplification

For *L. kluyveri* GO analyses, gene ontology information borrowed from *S. cerevisiae* (SGD, http://www.yeastgenome.org; GO Term Finder, version 0.86) was used to analyze the *L. kluyveri* genes. In brief, the *L. kluyveri* genes were converted to *S. cerevisiae* orthologs using a published Lk-to-Sc cross-reference table of orthology (Hsu et al. 2021). *L. kluyveri* genes in the input list without a clear *S. cerevisiae* ortholog (∼10%) were omitted from our GO enrichment analyses of “biological processes”. The enriched GO terms with adjusted *p*-values < 0.01 were considered significant (a more stringent cut-off for *p*-values generally did not affect the results of subsequent analyses). We did not correct for the genetic background because *L. kluyveri* and *S. cerevisiae* share similar gene numbers in their genomes (∼6000 genes). To browse the GO hierarchy, the enriched GO terms from each set of differentially expressed genes were placed on the QuickGo website (Binns et al. 2009). To “simplify” the enriched GO term list, the resolved GO terms were checked manually and the broader parent terms were removed, while the more specific child terms close to the leaf node of the hierarchy tree were retained to assist in simplifying functional annotations.

### Protein extraction and Western blotting

Total protein was extracted by using the NaOH/TCA/HU method with modifications (Lu et al. 2016). In brief, ∼3 OD_600_ cells were resuspended in 1 ml lysis buffer (0.185 M NaOH, 0.75% β-ME) and lysed on ice for 10 min. Next, we added 150 μl of 55 % trichloroacetic acid (TCA, T9159, Sigma, USA) to the lysis suspension at a final percentage of 8% to precipitate proteins. After 15 min on ice, the precipitants were spun down by centrifugation (Eppendorf 5424R centrifuge; FA-45-24-11 rotor; 13K rpm; 10 min; 4°C) and the supernatant was discarded. The centrifugation step was repeated to remove all residual supernatant. Then, we added 300 μl of high urea (HU) sample buffer (8 M Urea, 5% SDS, 0.2 M Tris-HCl pH 6.5, 1 mM Na_2_EDTA, 0.01% bromophenol blue) supplemented with 2M Tris base in a 50-to-3 ratio to the protein pellets, and then incubated them at 65°C for at least 40 min, before subjecting them to vigorous pipetting to sufficiently dissolve the pellets.

Protein extracts were resolved by 8% SDS-PAGE (Sambrook 2001) under a constant voltage of 80 V for 40 min of stacking followed by 150 V for 2 h of separating, before being blotted onto an Immobilon^®^-P PVDF Membrane (IPVH00010, Millipore by Merck) in a transfer buffer (25 mM Tris base, 192 mM glycine, 20% methanol) under a constant voltage of 30 V for 20 h of transfer at 4°C. Post-transfer membranes were then blocked with the blocking buffer [1% casein (C5890, Sigma) in PBST (1% Tween^®^ 20 in 1X PBS)] for 1 h. For immunodetection, rabbit polyclonal anti-TAP antibody (CAB1001, Thermo Fisher Scientific) diluted 1:2000 in blocking buffer was used to detect TAP-tagged Sef1 and Azf1. Alpha-tubulin was used as an internal control and it was detected by means of rabbit monoclonal antibody (ab184970, Abcam, UK) diluted 1:20000 in blocking buffer. The blots were hybridized with each primary antibody for 2 h at room temperature and washed three times with PBST (5 min each). Finally, the blot was hybridized with Peroxidase-AffiniPure Donkey Anti-Rabbit IgG secondary antibody (711-035-152, Jackson ImmunoResearch, USA) diluted 1:20000 in blocking buffer for 1 h at room temperature, and then washed three times with PBST. Chemiluminescence was developed with enhanced ECL reagents (NEL105001EA, PerkinElmer, USA) and visualized by exposure to an X-ray film.

### Statistical analysis

Details of statistical analyses are presented in the main text or corresponding figure legends. Statistical significance was established using unpaired Student’s t-tests in Excel 2016 or the default functions packaged in each analysis tool.

### Data reproducibility and replication

For all quantitative assays, the means were calculated from at least 3 technical repeats. For all qualitative assays, the experiments were repeated at least twice and one consistent and representative result was shown.

### Data availability

The RNA-seq data have been submitted to the Gene Expression Omnibus database (http://www.ncbi.nlm.nih.gov/geo/) under the accession number GSE200532. The processed results are shown in Table S1, S2, and S5–S12. The DNA-seq data have been submitted to the NCBI BioProject database (https://www.ncbi.nlm.nih.gov/bioproject) under the accession number PRJNA825358. The processed results are shown in Table S4.

## Acknowledgments

We are grateful to Professor Kenneth H. Wolfe (UCD, Ireland) for generously providing the DNA sequences of the *L. kluyveri* MAT loci used in this study. We are thankful to Dr. Shu-Yun Tung, the Genomics Core Laboratory (IMB, Academia Sinica, Taiwan) for technical assistance with the RNA-seq experiments. We thank all the JYL lab members for helpful discussions and comments on the manuscript. We thank John O’Brien for manuscript editing.

This work was supported by Academia Sinica of Taiwan (grant no. AS-IA-110-L01 and AS-GCS-110-01, to JYL) and the Taiwan Ministry of Science and Technology (grant no. MOST 110-2326-B-001-007, to JYL). PCH was supported by a MOST postdoctoral fellowship (MOST 110-2811-B-001-581). The funders had no role in study design, data collection and analysis, decision to publish, or preparation of the manuscript.

## Author contributions

Conceptualization: P.-C.H., J.-Y.L.; Methodology: P.-C.H., Y.-H.C., C.-W.L.; Validation: P.-C.H.; Formal analysis: P.-C.H., Y.-H.C., C.-W.L.; Investigation: P.-C.H., Y.-T.J., F.J.G.O., A.A.A.A., J.-Y.L.; Resources: J.-Y.L.; Data curation: P.-C.H., Y.-H.C., C.-W.L.; Writing – original draft: P.- C.H., J.-Y.L.; Writing – review & editing: P.-C.H., J.-Y.L.; Visualization: P.-C.H.; Supervision: J.-Y.L.; Project administration: J.-Y.L.; Funding acquisition: J.-Y.L.

## Conflicts of interest

The authors declare that they have no conflict of interest.

## Source data

**Figure 3-source data-1.** (1) Raw images for Figure 3D. Top and bottom images are X-ray films exposed to the same blot for 15 and 30 sec, respectively. (2) Raw images with labels for Figure 3D.

**Figure 3-source data-2.** (1) Raw images for Figure 3I. Top and bottom images are X-ray films exposed to the same blot for 30 sec and 5 min, respectively. (2) Raw images with labels for Figure 3I.

## Supplementary figures

**Figure S1.**
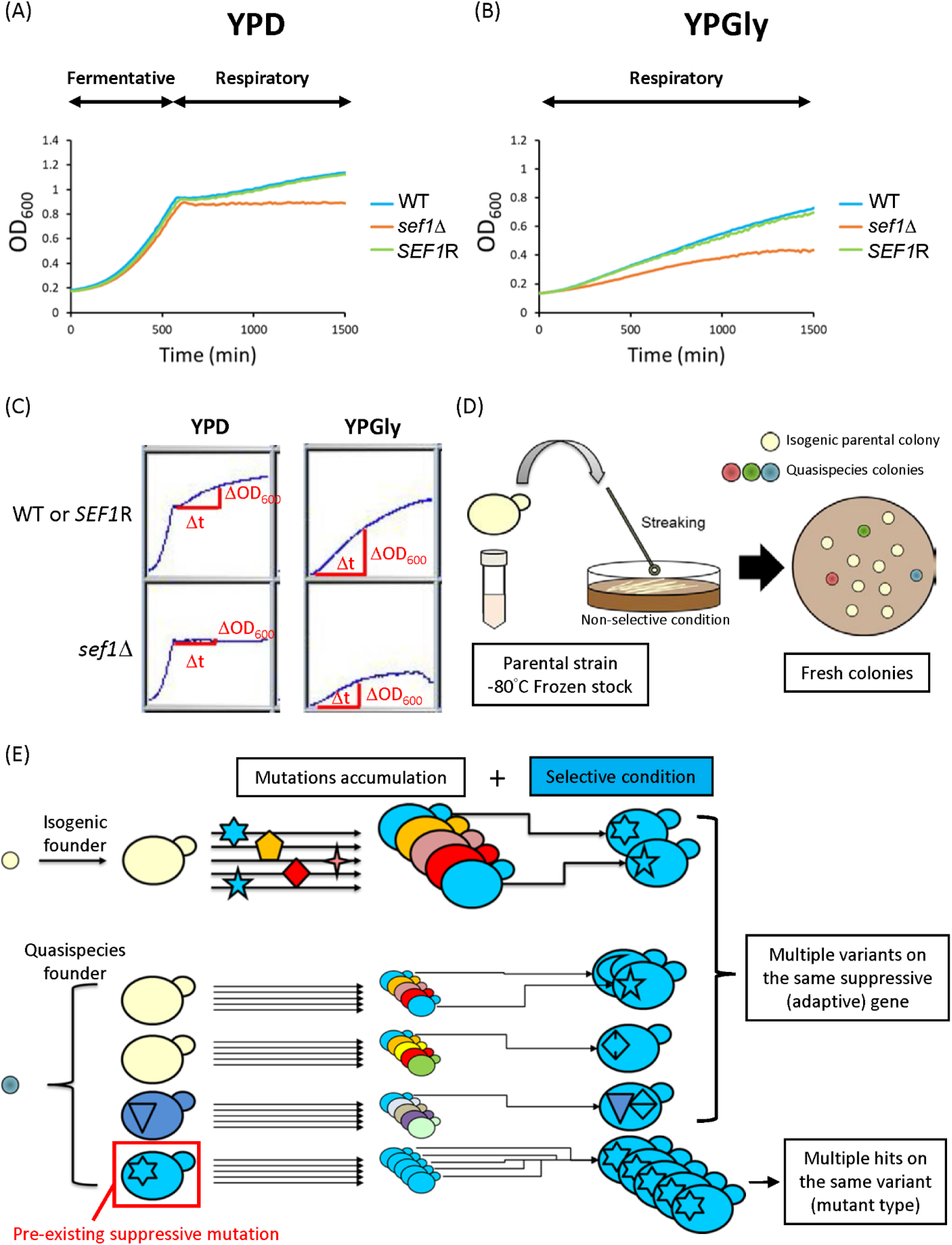
Deletion of *SEF1* affects fitness and the potential trajectories of adaptive evolution. (A) Growth curves of the *sef1*Δ mutant in YPD at 28°C. (B) Growth curves of the *sef1*Δ mutant in YPGly at 28°C. (C) Schematic representation of the maximal slope growth rate calculation for the post-diauxic shift growth phase in YPD and log-phase growth in YPGly. (D) Schematic representation of possible pre-existing genetic variations in the genomes of different individuals in founder colonies. (E) Schematic representation of suppressor formation by selection on pre-existing variations of a quasispecies founder or on new (*de novo*) adaptive mutations.

**Figure S2.**
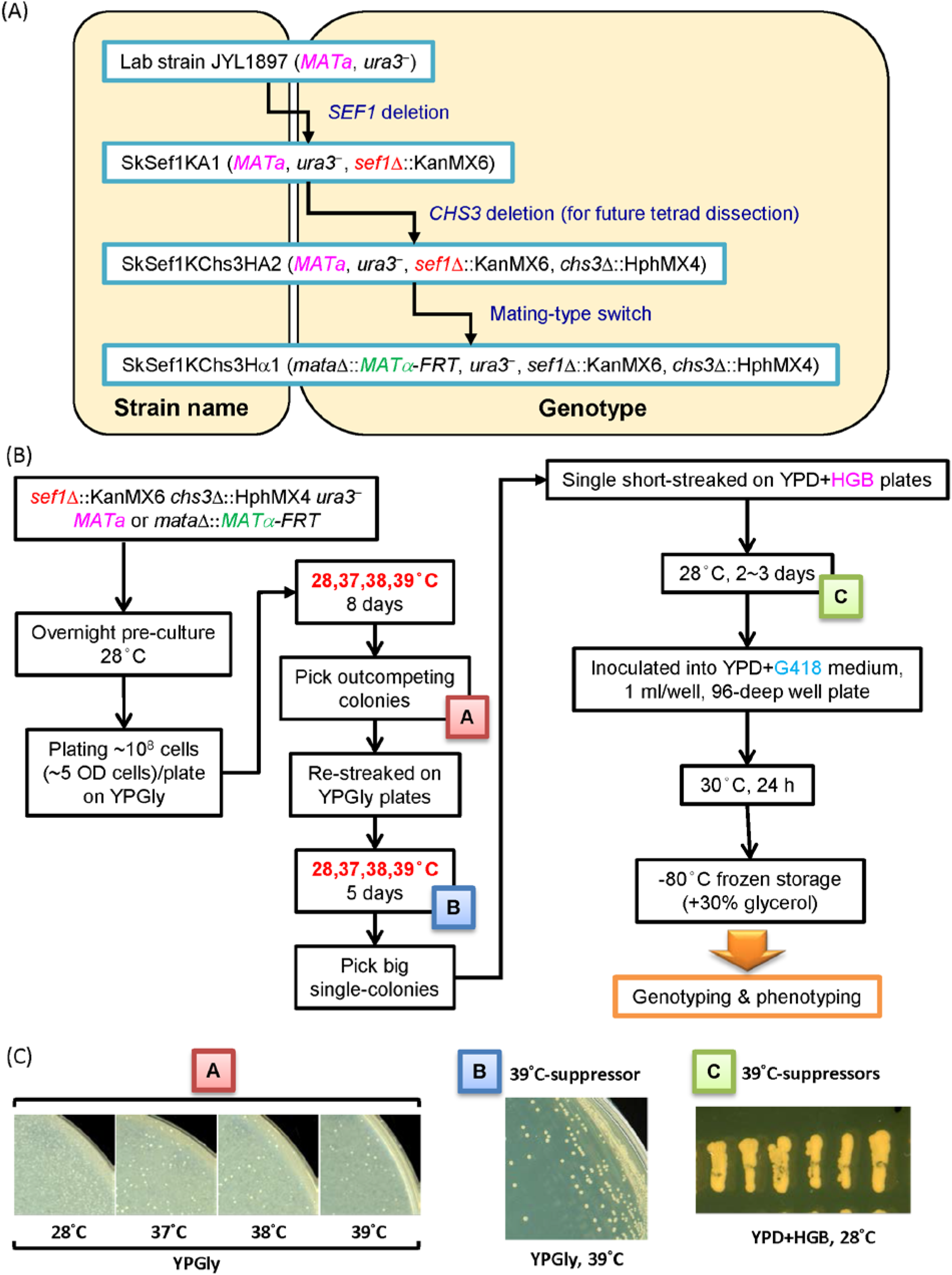
Workflow of *sef1*Δ suppressor development. (A) Construction of the *sef1*Δ founder strain. (B) Procedures for *sef1*Δ suppressor development and selection. (C) Examples of suppressor clone picking and purification steps.

**Figure S3.**
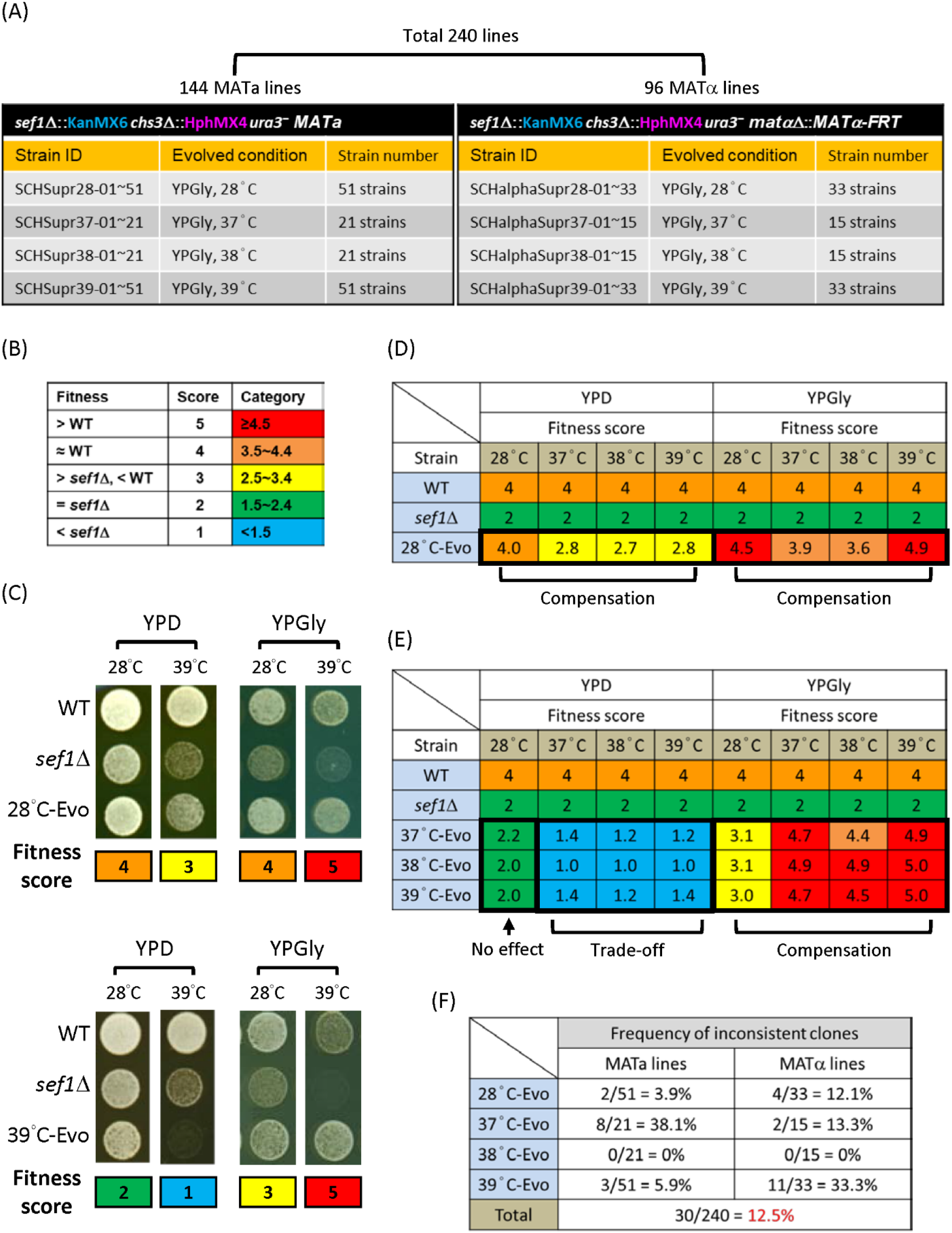
Summary of 240 *sef1*Δ suppressors. (A) Descriptions of all suppressors. (B) Criteria for simple fitness scoring and color-specified categories of *sef1*Δ suppressors. (C) Examples of growth phenotypes and simple fitness scores. (D) Mean simple fitness score of all 28°C-Evo *sef1*Δ suppressors. (E) Mean simple fitness score of all 37°C-, 38°C-, or 39°C-Evo *sef1*Δ suppressors. (F) Frequency of phenotypically inconsistent suppressor clones.

**Figure S4.**
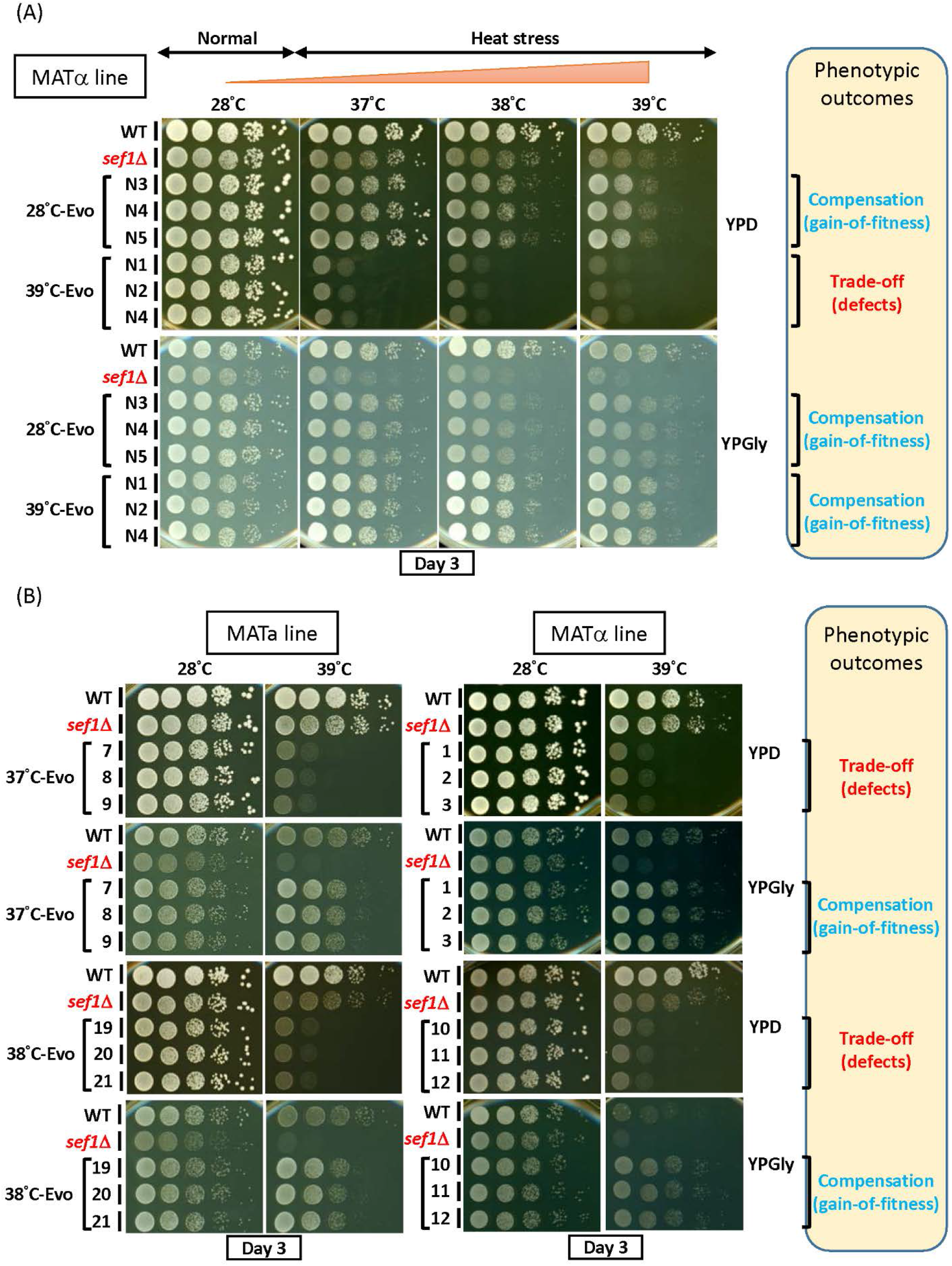
Phenotypic verification of *sef1*Δ suppressors with consistent phenotypes. (A) The suppressive growth phenotypes of re-purified *sef1*Δ suppressor clones (28°C-Evo and 39°C-Evo, MATα line). (B) The suppressive growth phenotypes of other randomly selected *sef1*Δ suppressor clones (37°C-Evo and 38°C-Evo, both MATa and MATα lines).

**Figure S5.**
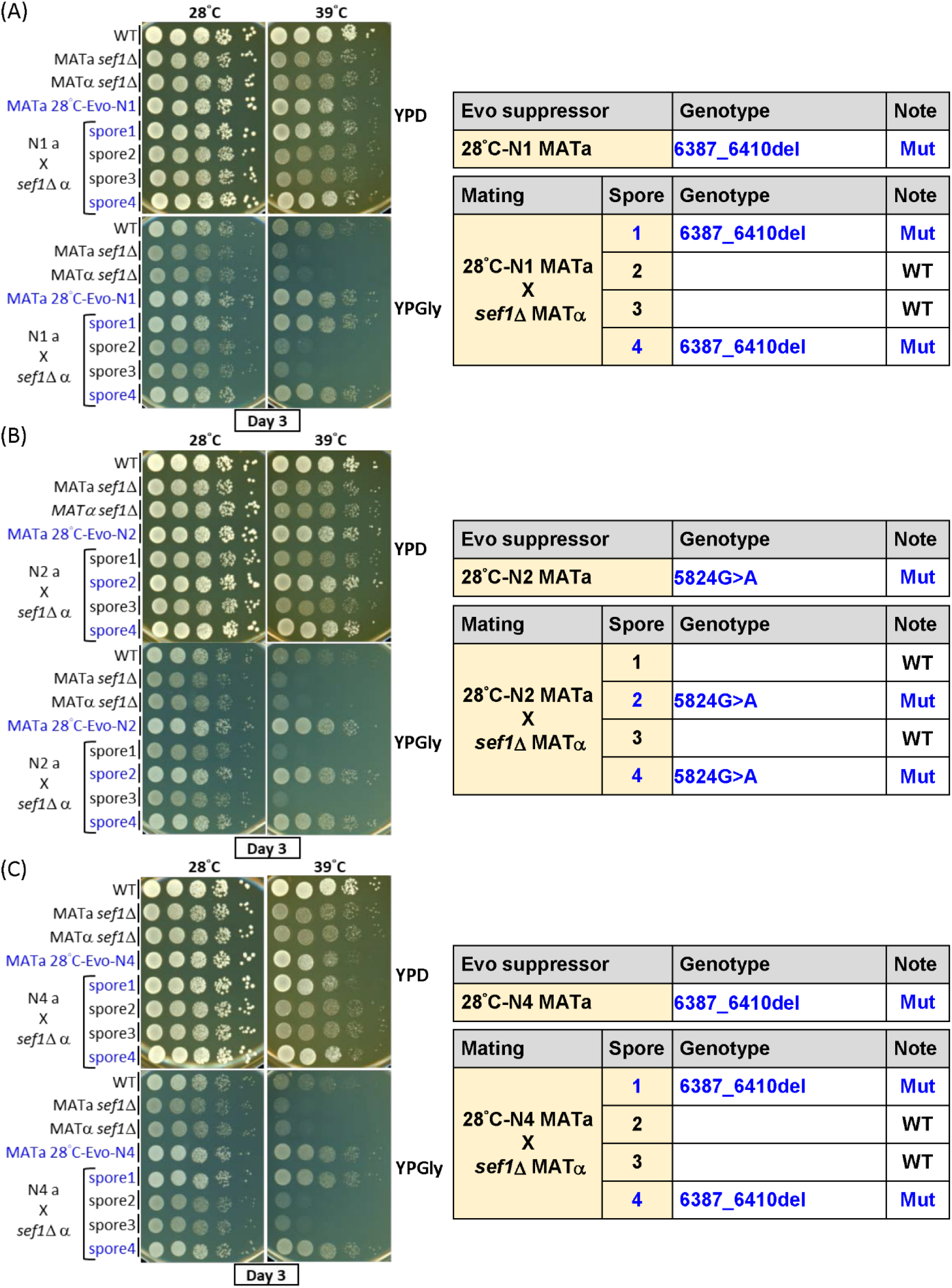
Genetic dissection of candidate causal mutations in MATa 28°C-Evo *sef1*Δ suppressors. Three clones (A) 28°C-Evo-N1, (B) 28°C-Evo-N2, and (C) 28°C-Evo-N4 were dissected. The genotypes of spores from each tetrad were checked by Sanger sequencing. Mut – mutant.

**Figure S6.**
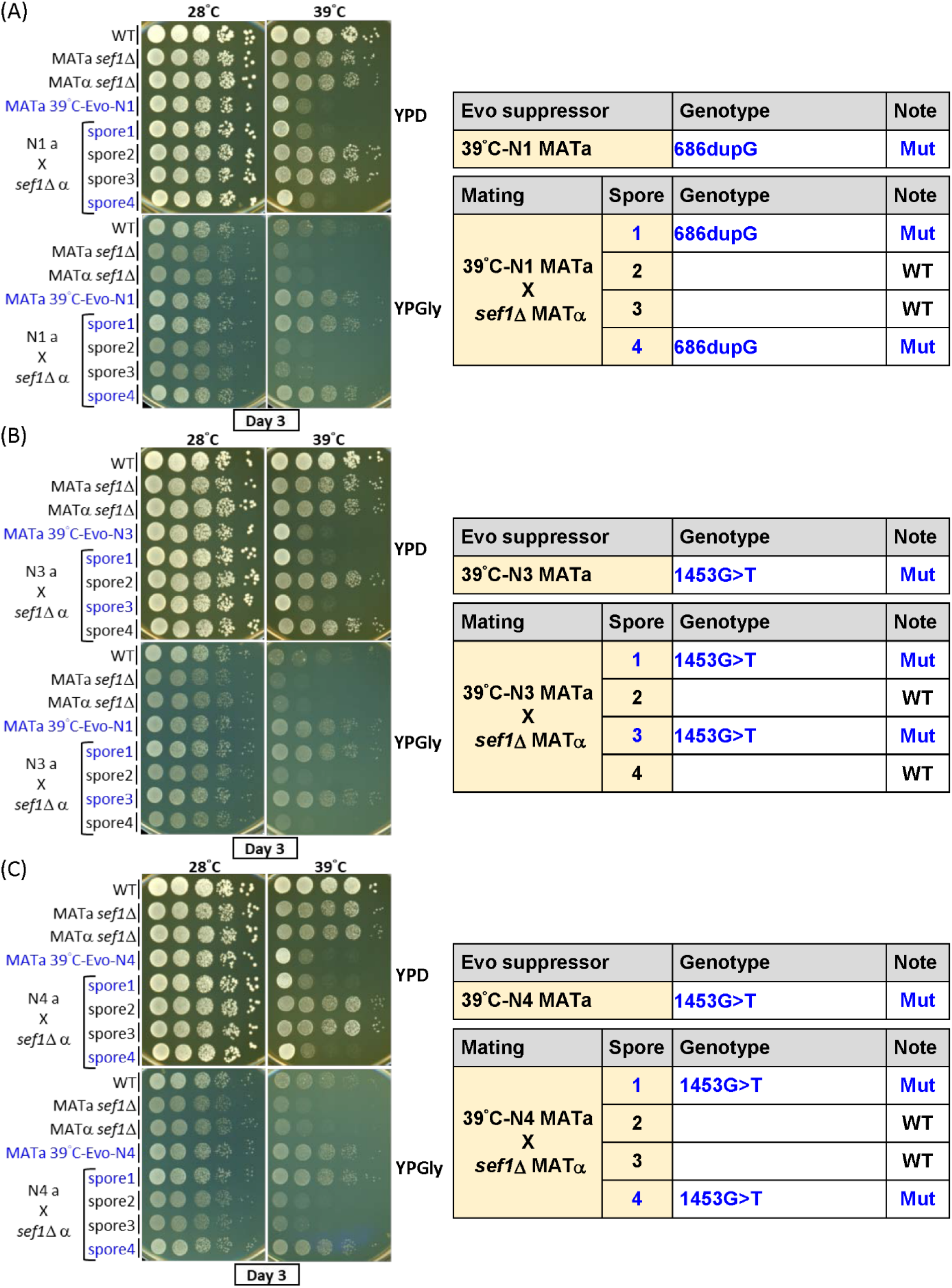
Genetic dissection of candidate causal mutations in MATa 39°C-Evo *sef1*Δ suppressors. Three clones (A) 39°C-Evo-N1, (B) 39°C-Evo-N3, and (C) 39°C-Evo-N4 were dissected. The genotypes of spores from each tetrad were checked by Sanger sequencing. Mut – mutant.

**Figure S7.**
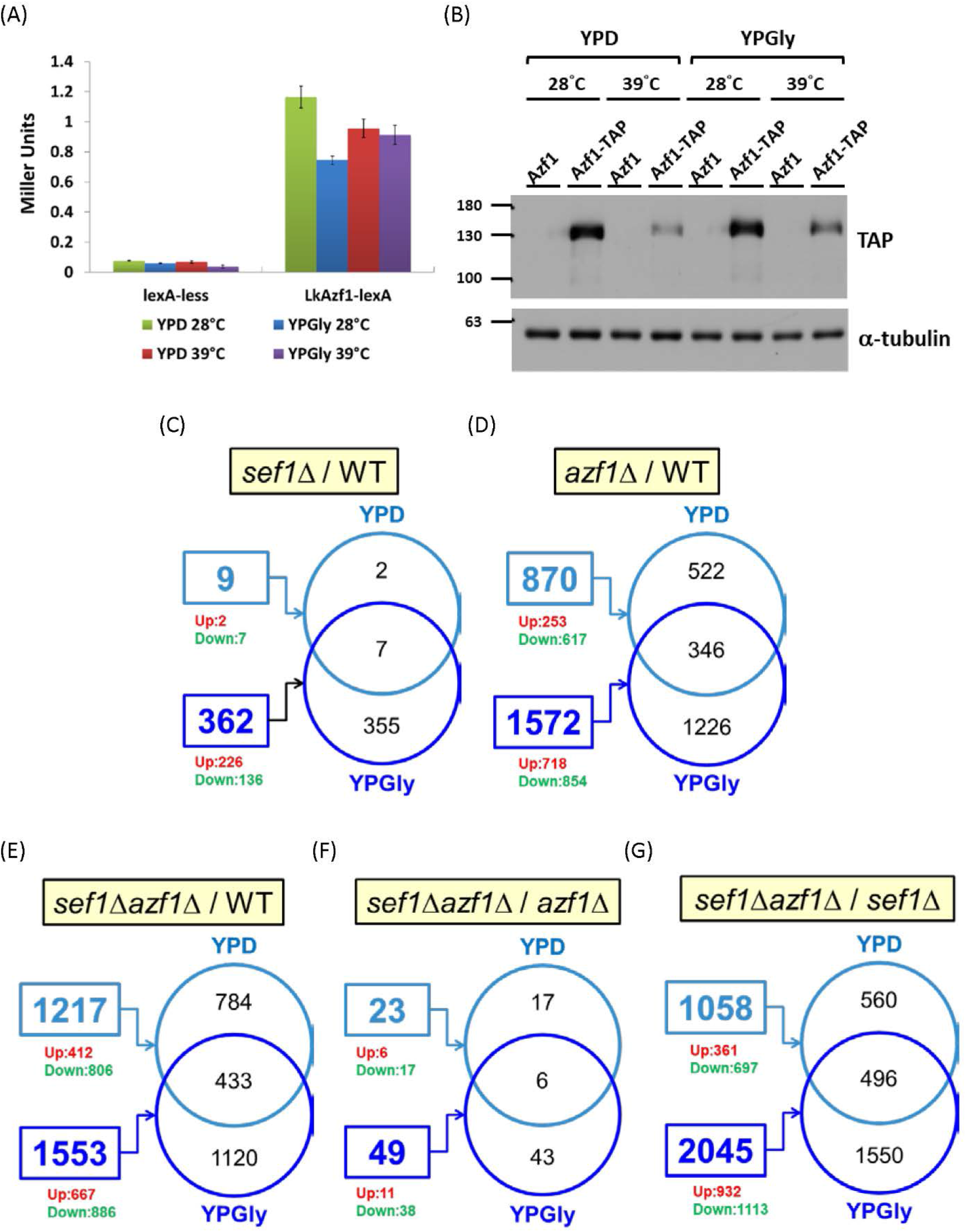
Differential gene expression in response to *azf1*Δ and *sef1*Δ mutations. (A) Heat stress (39°C) slightly reduces the transcriptional activation capability of Azf1, which was measured by one-hybrid assays. LacZ activity was measured by liquid-galactosidase assay and results are displayed as average Miller units ± SD from at least three technical repeats. (B) Azf1 protein abundance is reduced by heat stress (39°C). (C to G) Summaries of numbers of differentially expressed genes in *sef1*Δ/WT (C), *azf1*Δ/WT (D), *sef1*Δ*azf1*Δ/WT (E), *sef1*Δ*azf1*Δ/*sef1*Δ (F), and *sef1*Δ*azf1*Δ/*azf1*Δ (G). Numbers in rectangles are the total numbers of differentially expressed genes under a specific condition. Up or Down: the numbers of upregulated or downregulated genes, respectively. Venn diagrams display numbers of overlapping genes between the two conditions.

**Figure S8.**
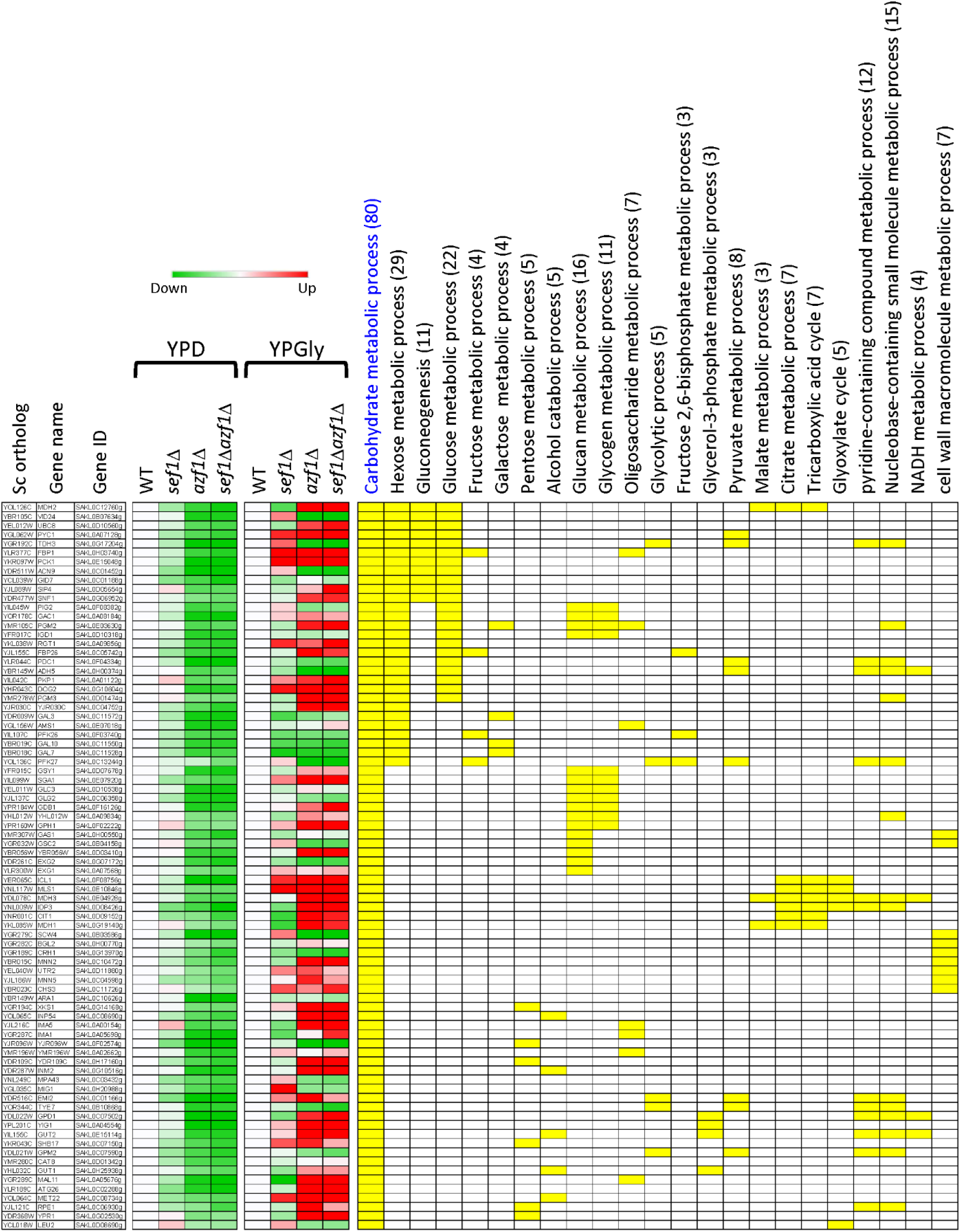
Dissection of downregulated carbohydrate metabolic process genes in response to *azf1*Δ mutation under the YPD condition. The heatmap was generated using the mean TPM ratio from RNA-seq data relative to the wild-type under each condition. The yellow blocks highlight the sub-GO groups to which each gene belongs. Total gene numbers for each GO group are specified in parentheses.

**Figure S9.**
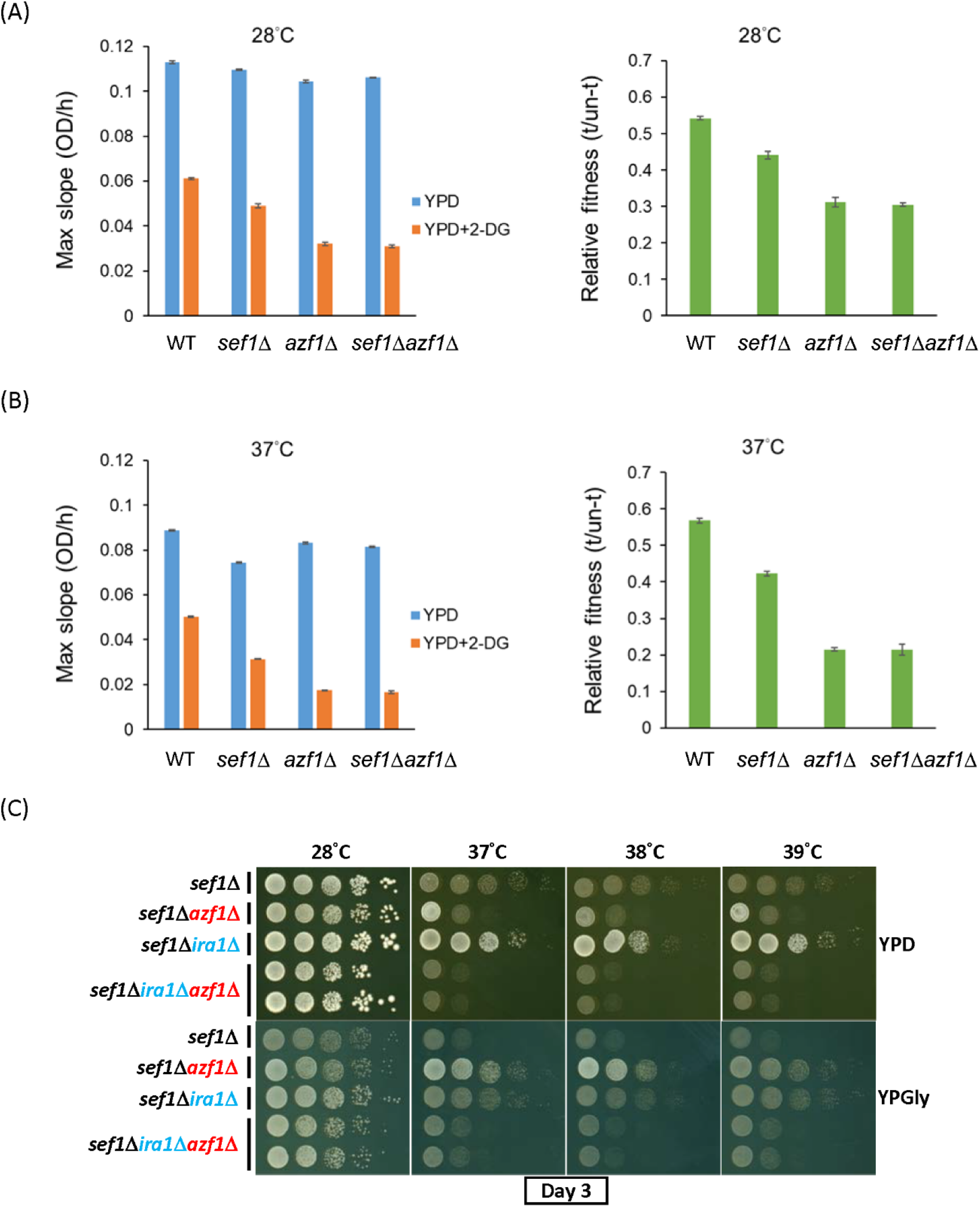
The fitness of *azf1*Δ cells in response to 2-deoxyglucose under the YPD condition. (A) Max slope growth rate and relative fitness of the *azf1*Δ mutants at 28°C. (B) Max slope growth rate and relative fitness of the *azf1*Δ mutants at 37°C. (C) Synthetic growth defect of *azf1*Δ*ira1*Δ in the *sef1*Δ background under heat-stressed conditions.

**Figure S10.**
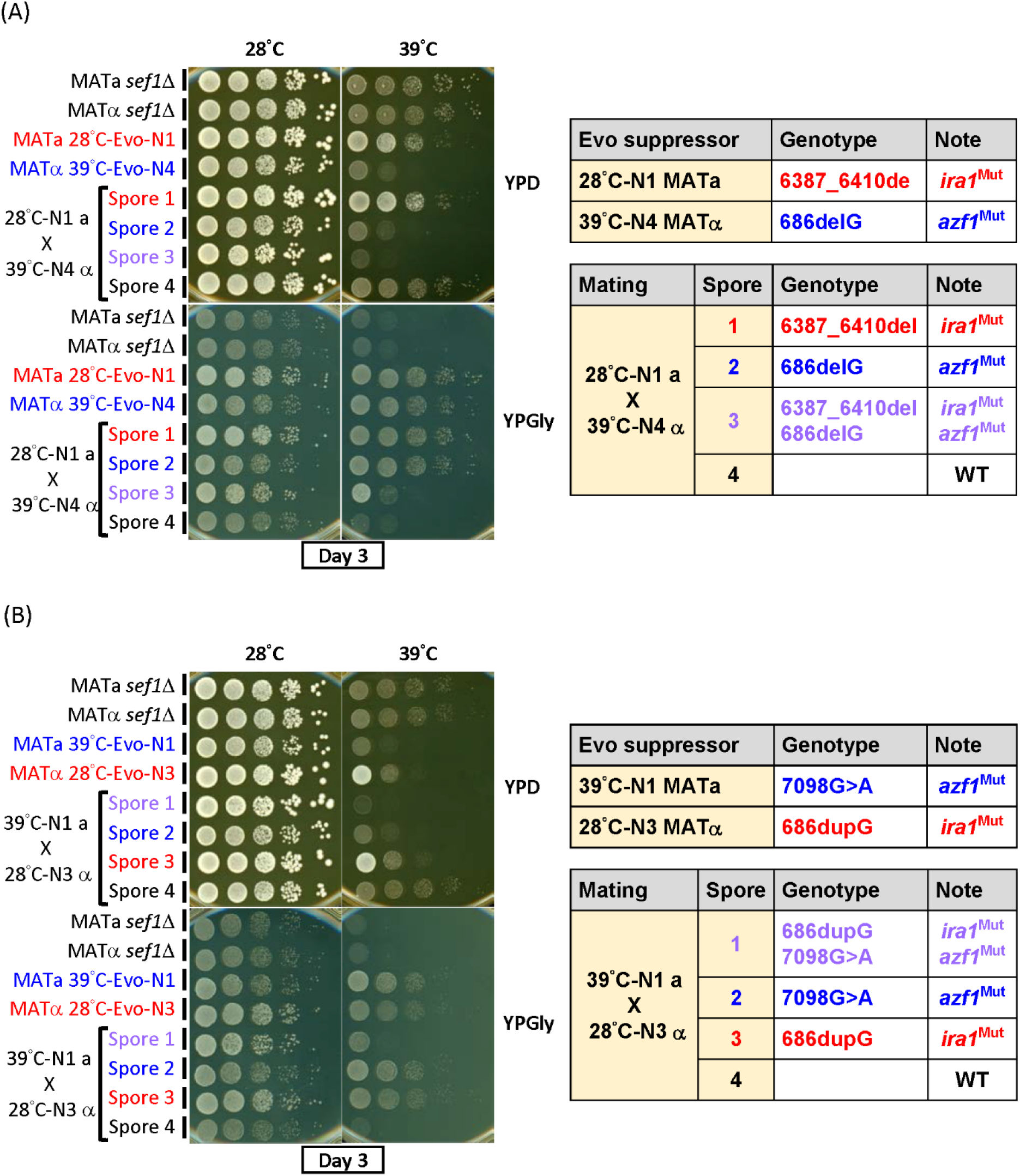
Synthetic effects of the *ira1* mutation from 28°C-Evo *sef1*Δ suppressors and the *azf1* mutation from 39°C-Evo *sef1*Δ suppressors. (A) Tetrad dissection and Sanger sequencing of 28°C-Evo-N1 MATa and 39°C-Evo-N4 MATα mating products. (B) Tetrad dissection and Sanger sequencing of 39°C-Evo-N1 MATa and 28°C-Evo-N3 MATα mating products. All four spores from each tetrad were phenotyped by using the spot assay. The *IRA1* and *AZF1* loci were sequenced.

**Figure S11.**
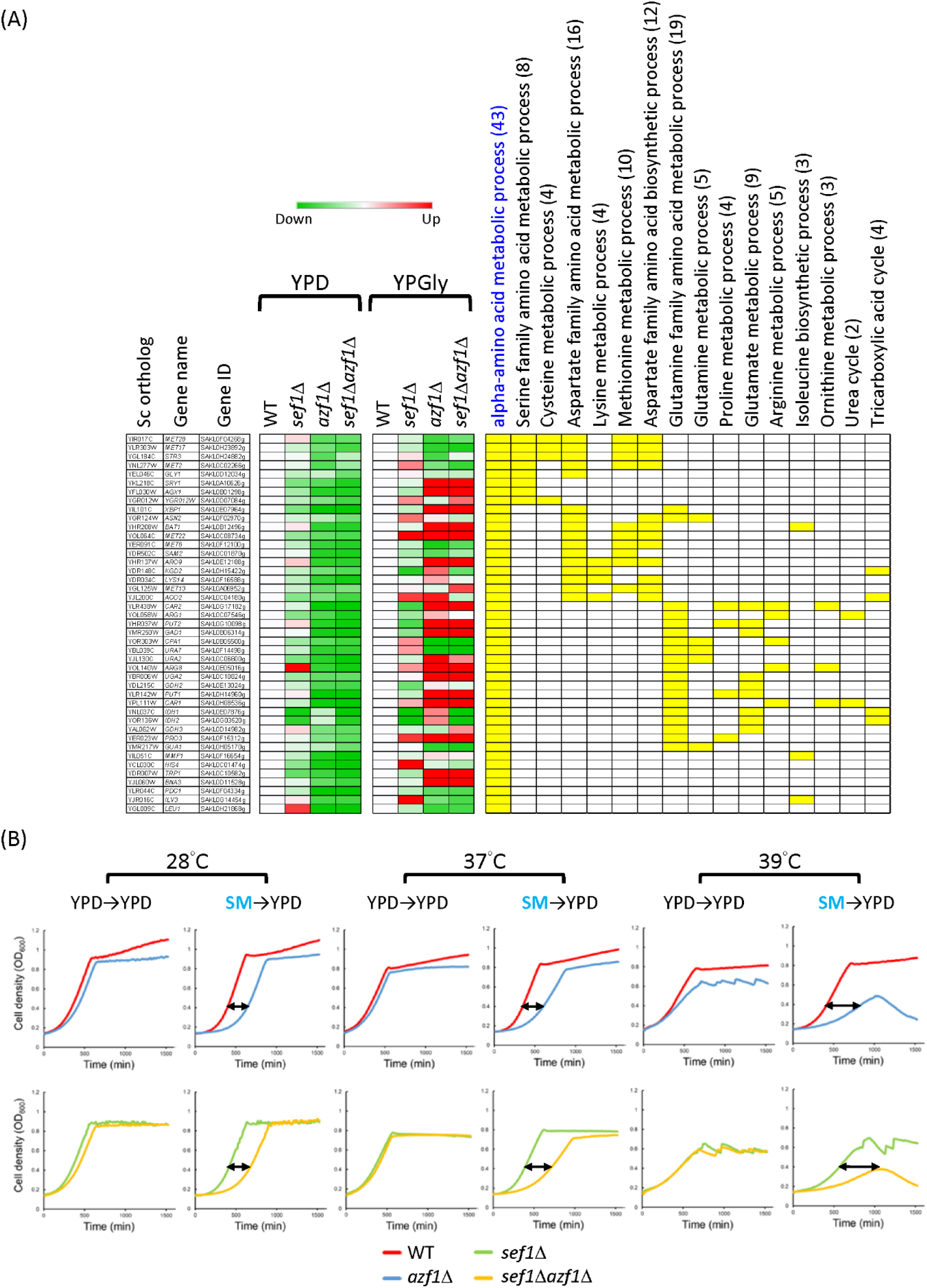
Dissection of downregulated alpha-amino acid metabolic process genes in response to *azf1*Δ mutation under the YPD condition. (A) The heatmap was generated using the mean TPM ratio from RNA-seq data relative to the wild-type under each condition. The yellow blocks highlight the sub-GO groups to which each gene belongs. Total gene numbers in each GO group are specified in parentheses. (B) The effect of pre-amino acid starvation (23-h starvation in SM+2X uracil medium) on the growth of the *azf1*Δ mutants at indicated temperatures. The jagged curves reflect cellular aggregation or the presence of dead cells mixed with live cells under harsher culture environments. The near-concave curves (39°C, SM to YPD curves) were caused by severe cell death.

**Figure S12.**
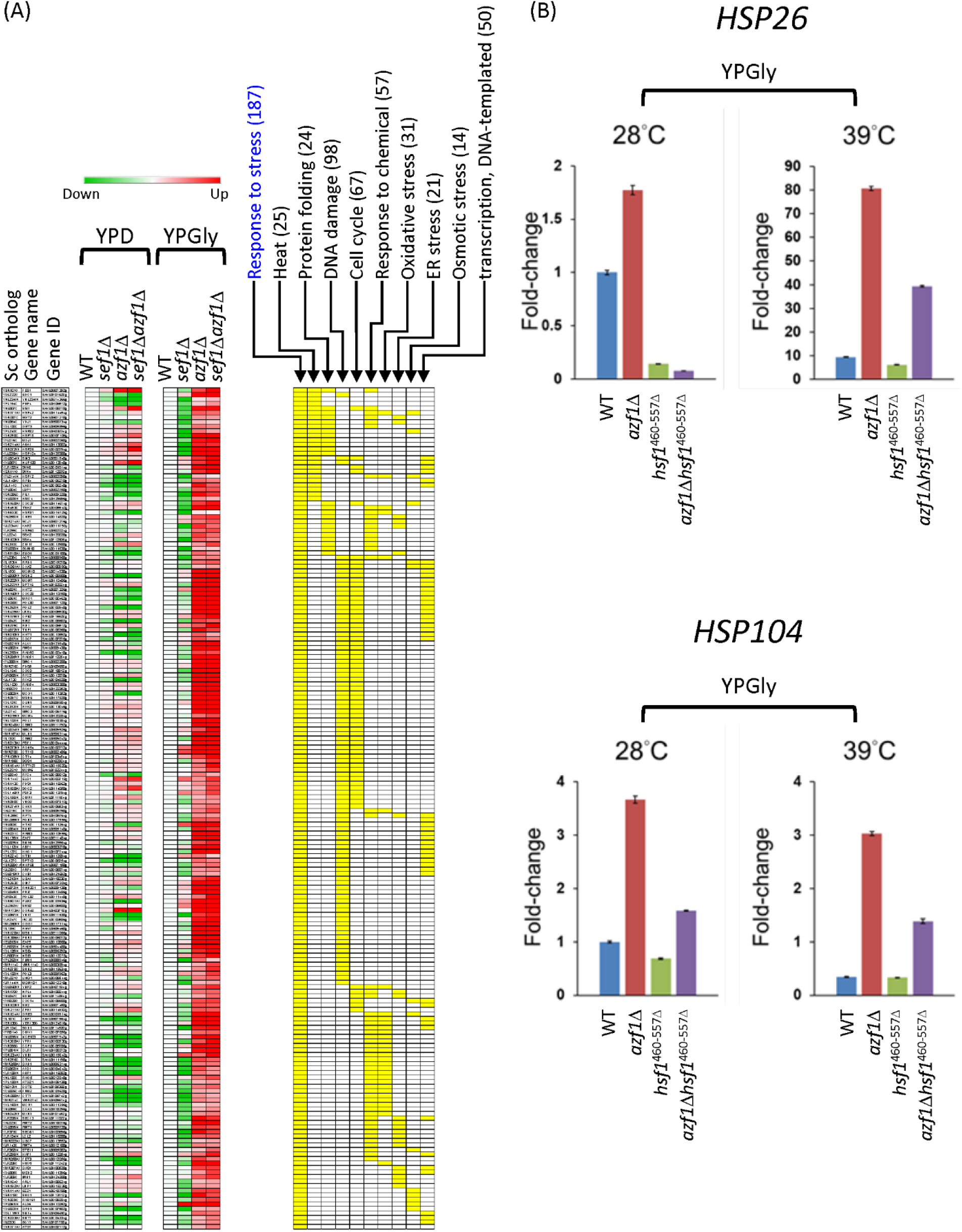
Dissection of upregulated stress response genes in response to *azf1*Δ mutation under the YPGly condition. (A) The heatmap was generated using the mean TPM ratio from RNA-seq data relative to the wild-type under each condition. The yellow blocks highlight the sub-GO groups to which each gene belongs. Total gene numbers in each GO group are specified in parentheses. (B) Expression of *HSP26* and *HSP104* in response to hypomorphic *hsf1* mutation (a truncated *hsf1* with the C-terminal 460-557 amino acids removed) under the YPGly condition. The relative fold-change of each gene is shown as 2^−ΔΔCT^, using *CDC34* (SAKL0D02530g) as the endogenous control and the ΔC_T_ value from the wild-type sample as the corresponding calibration value. Expression levels are displayed as means ± SD from three technical repeats.

**Figure S13.**
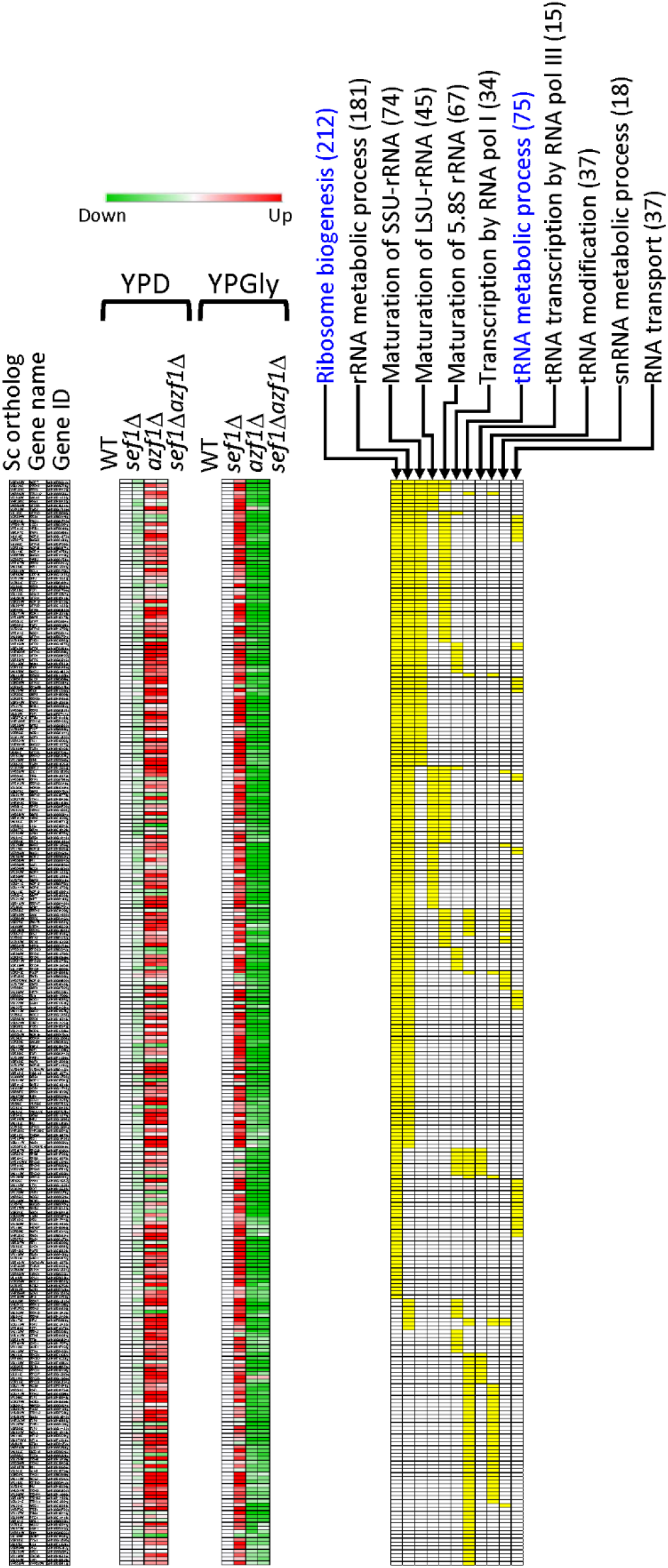
Dissection of the downregulated ribosome- and tRNA-related genes in response to *azf1*Δ mutation under the YPGly condition. The heatmap was generated using the mean TPM ratio from RNA-seq data relative to the wild-type under each condition. The yellow blocks highlight the sub-GO groups to which each gene belongs. Total gene numbers in each GO group are specified in parentheses.

**Figure S14.**
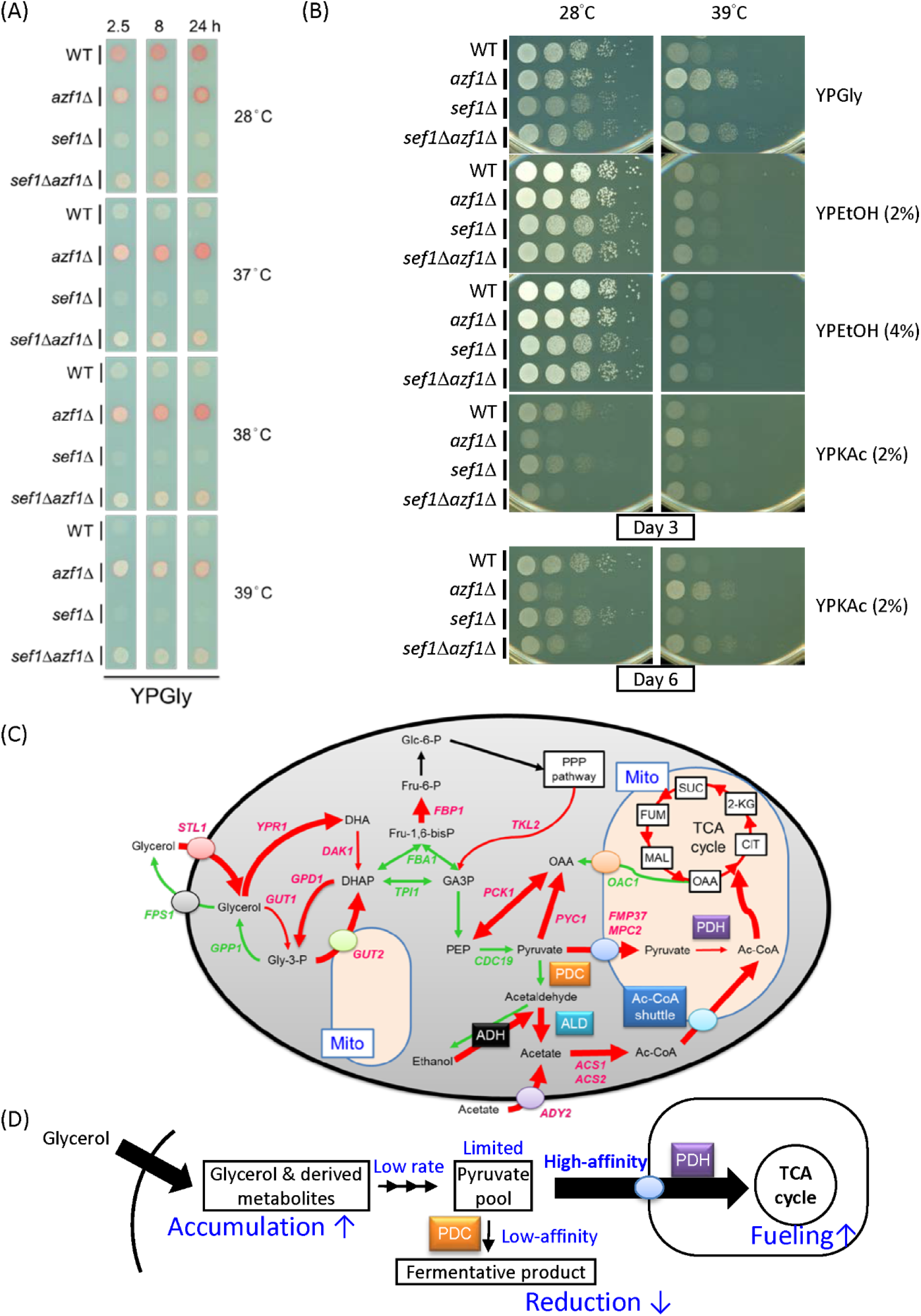
Glycerol and acetate, but not ethanol, are required for the enhanced fitness of *azf1*Δ mutants under heat-stressed conditions. (A) The *azf1*Δ mutants maintain relatively higher TTC reduction activity under the YPGly condition compared to *sef1*Δ strains. The formation of red product in the cell colonies indicates that the cells have competent TTC reduction activity. The whiter spots indicate defects in cellular respiration. (B) Acetate, but not ethanol, endows weaker heat resistance on the *azf1*Δ mutants. YPEtOH (YP + ethanol); YPKAc (YP + potassium acetate). Concentrations of ethanol and acetate are shown in parentheses. (C) Remodeled glycerol utilization in *azf1*Δ cells. Gly-3-P: glycerol-3-phosphate; DHA: dihydroxyacetone; DHAP: dihydroxyacetone phosphate; GA3P: glyceraldehyde-3-phosphate; Glc-6-P: glucose-6-phosphate; Fru-6-P: fructose-6-phosphate; Fru-1,6-bisP: fructose-1,6-bisphosphate; PPP: pentose phosphate pathway; PEP: phosphoenolpyruvate; Ac-CoA; acetyl coenzyme A; PDC: pyruvate decarboxylase complex; PDH: pyruvate dehydrogenase complex; ALD: aldehyde dehydrogenase; ADH: alcohol dehydrogenase; OAA: oxaloacetate; CIT: citrate; 2-KG; 2-oxoglutarate; SUC: succinate; MAL: malate. Mito: mitochondrion. Red arrow: upregulated gene; green arrow: downregulated gene. The thickness of the arrows reflects the relative RNA abundance according to the heatmap presented in Figure 5D. (D) Proposed glycerol-driven metabolic remodeling at the pyruvate node in *azf1*Δ cells. In this model, glycerol accumulates intracellularly due to enhanced uptake, but it is converted to pyruvate at a low rate to maintain a limited pyruvate pool. Consequently, high-affinity mitochondrial pyruvate carriers plus PDH complex compete for the limited pyruvate with the low-affinity PDC complex, thereby fueling respiration rather than fermentation. Accordingly, *azf1*Δ cells benefit from the mitochondrial activity, supporting survival upon encountering heat stress.

**Figure S15.**
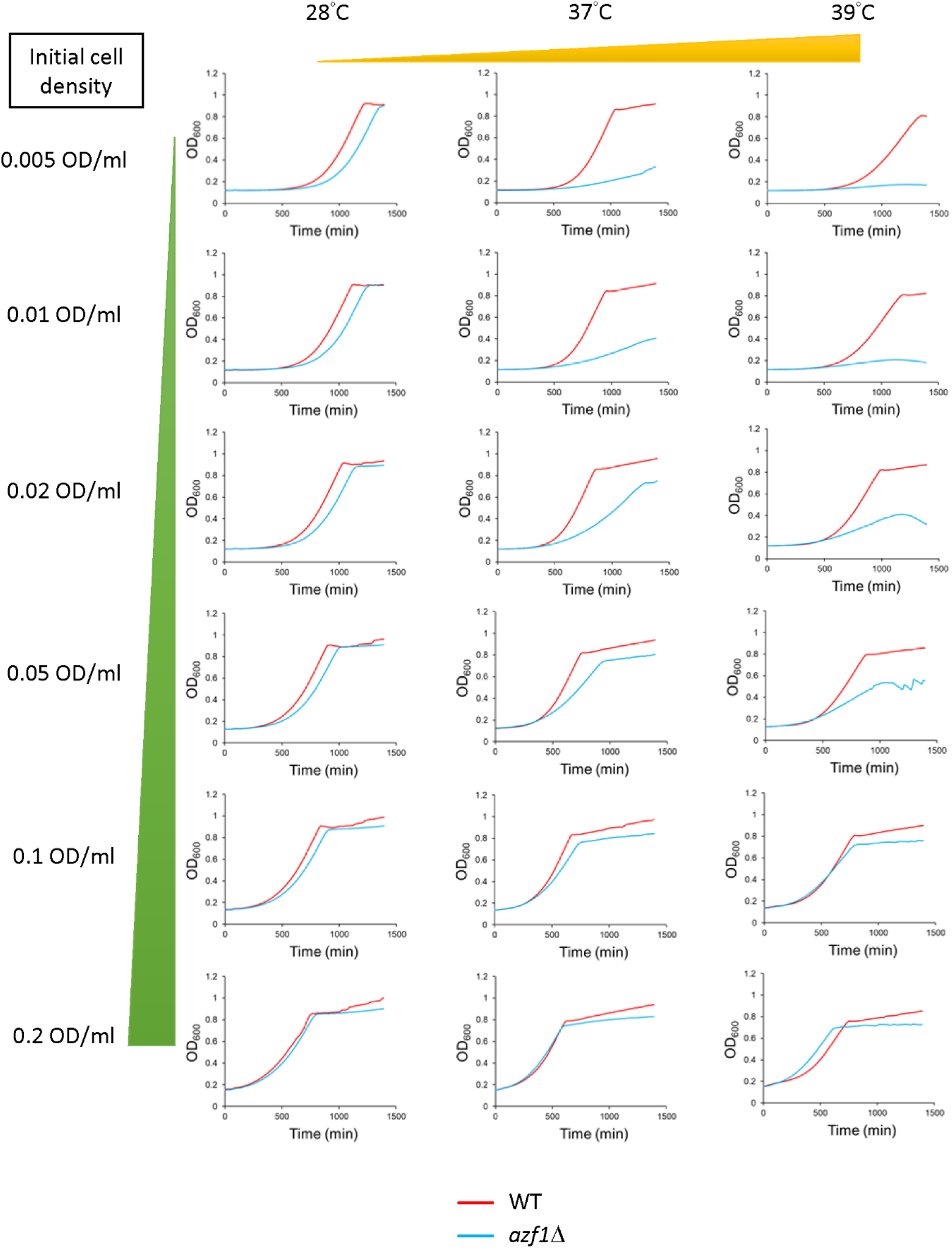
Growth curves of wild-type and *azf1*Δ mutant cells in YPD in response to increasing initial inoculum densities and temperature. Representative source data for Figure 6A.

**Figure S16.**
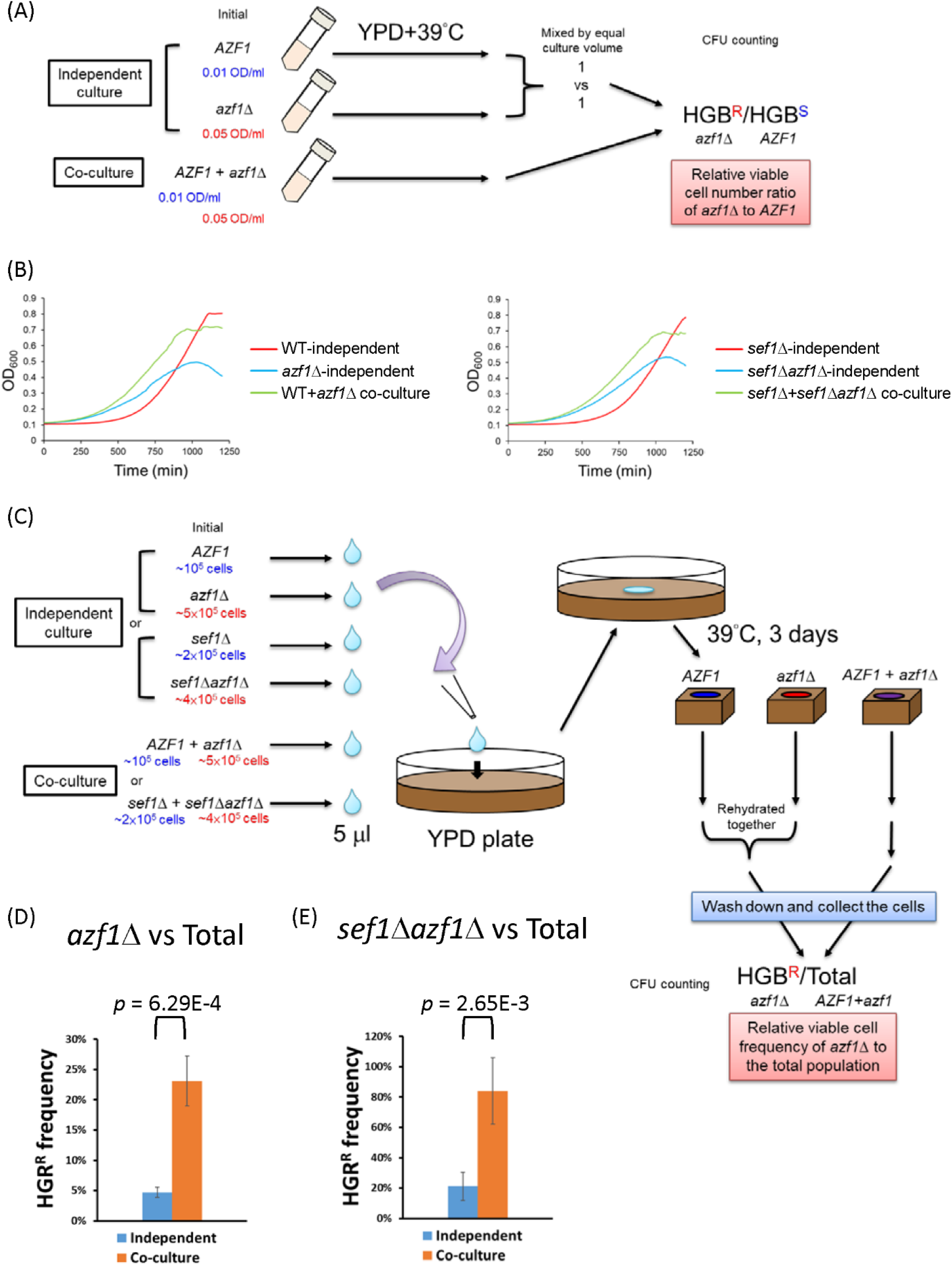
Cooperative growth assays on the *AZF1* and *azf1*Δ strains. (A) Illustrative workflow of the cooperative growth assay on the *AZF1* and *azf1*Δ strains in YPD liquid broth at 39°C. Growth in a 96-well plate was measured on a Tecan plate reader with intermittent shaking. Colony-forming units (CFUs) were counted by plating on YPD (total) and then replicated to a YPD+HGB plate to distinguish HGB-resistant *azf1*Δ strains and HGB-sensitive *AZF1* strains. (B) Source growth curves of Figure 6B and 6C. (C) Illustrative workflow of the cooperative growth assay on *AZF1* and *azf1*Δ strains on a YPD plate at 39°C. (D) The *azf1*Δ cells proved more persistent when co-grown with wild-type cells on an agar plate under the “Dex-trade-off” condition. (E) The *sef1*Δ*azf1*Δ cells proved more persistent when co-grown with *azf1*Δ cells on an agar plate under the “Dex-trade-off” condition. For (D) and (E), statistical significance tests were carried out using unpaired Student’s t-tests.

**Figure S17.**
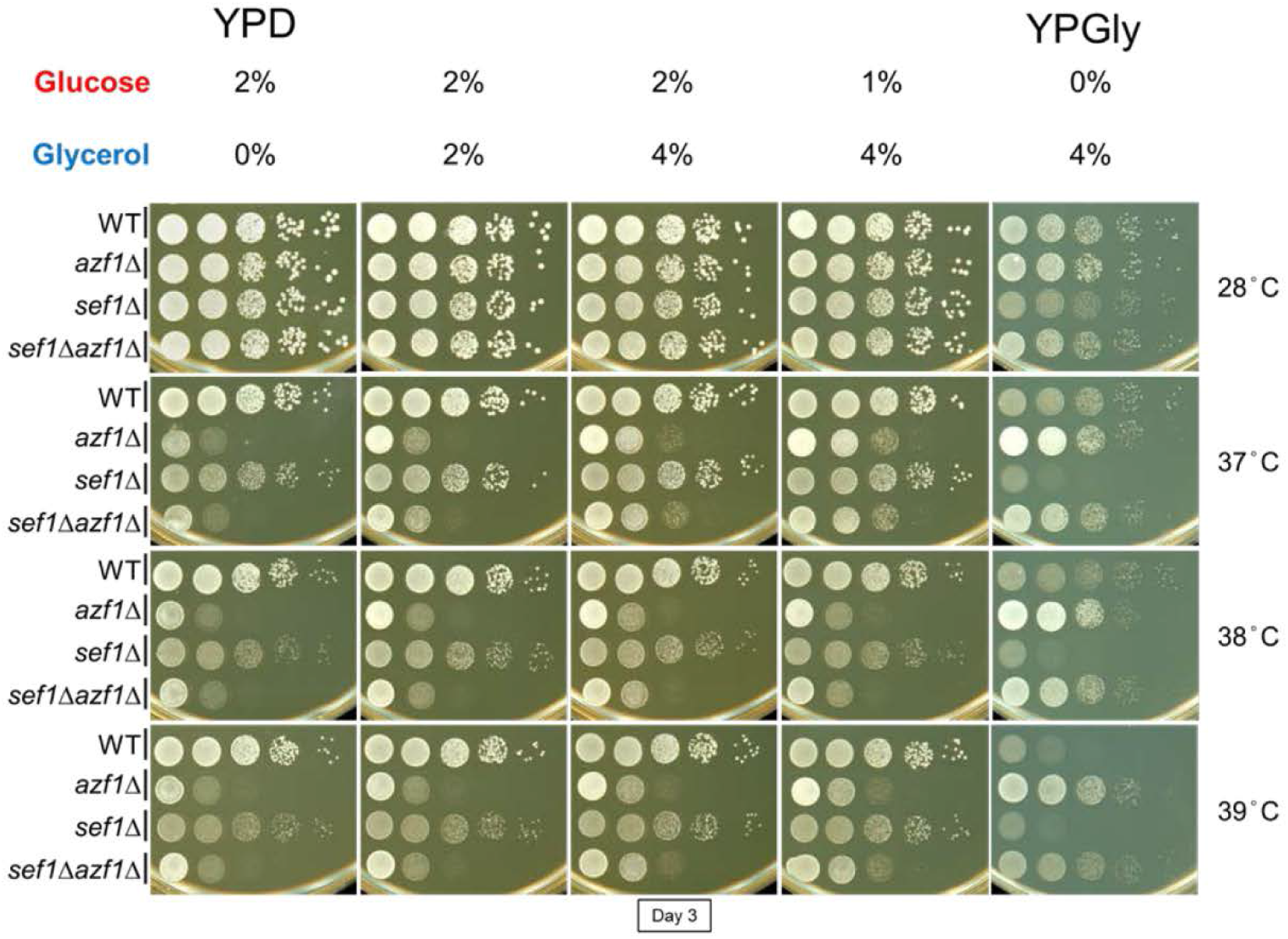
Effects of mixed glucose and glycerol on growth of *azf1*Δ cells. Increasing the glycerol concentration in YPD did not drastically ameliorate the “Dex-trade-off” effect. The *azf1*Δ mutants grew slightly better in YPD+4%Gly than in YPD, but still clearly worse than in YPGly. This outcome is possibly due to the protective effect of the elevated osmolarity generated by 4% glycerol.

## Supplementary tables

**Table S1. DESeq2 results of *sef1*Δ vs. wild-type gene expression profiles in YPD.**

**Table S2. DESeq2 results of *sef1*Δ vs. wild-type gene expression profiles in YPGly.**

**Table S3. Strain list for 240 *sef1*Δ suppressors and simple fitness scores.**

**Table S4. Mutation spectra of 12 sequenced *sef1*Δ suppressors.**

**Table S5. DESeq2 results of *azf1*Δ vs. wild-type gene expression profiles in YPD.**

**Table S6. DESeq2 results of *azf1*Δ vs. wild-type gene expression profiles in YPGly.**

**Table S7. DESeq2 results of *sef1*Δ*azf1*Δ vs. wild-type gene expression profiles in YPD.**

**Table S8. DESeq2 results of *sef1*Δ*azf1*Δ vs. wild-type gene expression profiles in YPGly.**

**Table S9. DESeq2 results of *sef1*Δ*azf1*Δ vs. *azf1*Δ gene expression profiles in YPD.**

**Table S10. DESeq2 results of *sef1*Δ*azf1*Δ vs. *azf1*Δ gene expression profiles in YPGly.**

**Table S11. DESeq2 results of *sef1*Δ*azf1*Δ vs. *sef1*Δ gene expression profiles in YPD.**

**Table S12. DESeq2 results of *sef1*Δ*azf1*Δ vs. *sef1*Δ gene expression profiles in YPGly.**

**Table S13. Strains, primers, plasmids, media, and important chemicals.**

## Supplementary source data

**Figure S3-source data 1.** All spot assay results for 240 *sef1*Δ suppressors.

**Figure S7-source data-1.** (1) Raw images Figure S7B. Top and bottom images are X-ray films exposed to the same blot for 30 sec and 5 min, respectively. (2) Raw images with labels for Figure S7B.

## Notes

### Competing Interest Statement

The authors have declared no competing interest.

